# Disentangling top-down drivers of mortality underlying diel population dynamics of *Prochlorococcus* in the North Pacific Subtropical Gyre

**DOI:** 10.1101/2021.06.15.448546

**Authors:** Stephen J. Beckett, David Demory, Ashley R. Coenen, John R. Casey, Mathilde Dugenne, Christopher L. Follett, Paige Connell, Michael C.G. Carlson, Sarah K. Hu, Samuel T. Wilson, Daniel Muratore, Rogelio A. Rodriguez-Gonzalez, Shengyun Peng, Kevin W. Becker, Daniel R. Mende, E. Virginia Armbrust, David A. Caron, Debbie Lindell, Angelicque E. White, François Ribalet, Joshua S. Weitz

**Affiliations:** School of Biological Sciences, Georgia Institute of Technology, Atlanta, GA, USA; Center for Microbial Dynamics and Infection, Georgia Institute of Technology, Atlanta, GA, USA; School of Physics, Georgia Institute of Technology, Atlanta, GA, USA; Daniel K. Inouye Center for Microbial Oceanography: Research and Education, University of Hawai’i at Mānoa, Honolulu, HI, USA; Department of Oceanography, University of Hawai’i at Mānoa, Honolulu, HI, USA; Department of Earth, Atmospheric, and Planetary Sciences, Massachusetts Institute of Technology, Boston, MA, USA; Department of Biological Sciences, University of Southern California, Los Angeles, CA, USA; Biology Department, San Diego Mesa College, San Diego, CA, USA; Faculty of Biology, Technion – Israel Institute of Technology, Haifa, Israel; Department of Marine Chemistry and Geochemistry, Woods Hole Oceanographic Institution, Woods Hole, MA, USA; School of Natural and Environmental Sciences, Newcastle University, Newcastle upon Tyne, UK; Interdisciplinary Graduate Program in Quantitative Biosciences, Georgia Institute of Technology, Atlanta, GA, USA; Santa Fe Institute, Santa Fe, NM, USA; Adobe, San Jose, CA, USA; GEOMAR Helmholtz Centre for Ocean Research, Kiel, Germany; Laboratory of Applied Evolutionary Biology, Department of Medical Microbiology, Academic Medical Centre, University of Amsterdam, Amsterdam, The Netherlands; School of Oceanography, University of Washington, Seattle, WA, USA; Institut de Biologie, École Normale Supérieure, Paris, France

**Author notes:** present address: Sorbone Université, CNRS, UMR 7232 Biologie Intégrative des Organismes Marins (BIOM), Observatoire Océanologique, Banyuls-sur-Mer, France. present address: Physical and Life Sciences Directorate, Lawrence Livermore National Laboratory, Livermore, CA, USA. present address: Sorbonne Université, CNRS, UMR 7093, Laboratoire d’Océanographie de Villefranche-sur-Mer (LOV), Villefranche-sur-Mer, France. present address: Department of Earth, Ocean and Ecological Sciences, University of Liverpool, Liverpool, UK. present address: Department of Oceanography, Texas A&M University, College Station, TX, USA. These authors contributed equally (Stephen J. Beckett, David Demory). **Authorship Statement** S.J.B., D.D. and J.S.W. designed the study, with contributions from all authors. J.R.C., P.C., M.C.G.C., S.T.W., D.C., D.L., F.R. contributed to data collection, sample processing, and data preparation. S.T.W. served as chief scientist for the research expedition. S.J.B., D.D., A.R.C., R.A.R.-G., S.P., F.R., J.S.W. contributed to coding and data analysis. S.J.B, D.D., and J.S.W. led the writing of the manuscript with contributions from all the authors. All authors approved the final manuscript. **Data availability** Code for ECLIP and performing Bayesian parameter inference was written and run in Julia, with plotting performed in MATLAB. Core code for running model simulations, analysis and plotting is archived on Zenodo: https://doi.org/10.5281/zenodo.7937657 (Beckett *et al*., 2023).

## Abstract

Photosynthesis fuels primary production at the base of marine food webs. Yet, in many surface ocean ecosystems, diel-driven primary production is tightly coupled to daily loss. This tight coupling raises the question: which top-down drivers predominate in maintaining persistently stable picocyanobacterial populations over longer time scales? Motivated by high-frequency surface water measurements taken in the North Pacific Subtropical Gyre (NPSG), we developed multitrophic models to investigate bottom-up and top-down mechanisms underlying the balanced control of *Prochlorococcus* populations. We find that incorporating photosynthetic growth with viral- and predator-induced mortality is sufficient to recapitulate daily oscillations of *Prochlorococcus* abundances with baseline community abundances. In doing so, we infer that grazers function as the primary top-down factor despite high standing viral particle densities while identifying the potential for light-dependent viral traits and non-canonical loss factors to shape the structure and function of marine microbial communities.

## 1 Introduction

Highly resolved surface ocean observations reveal repeatable daily changes in the abundance of ubiquitous picocyanobacteria at the base of the marine microbial food web, including *Prochlorococcus* and *Synechococcus* (Ribalet *et al*., 2015; Vaulot & Marie, 1999). Typically, picocyanobacteria decrease in abundance during the day and then increase overnight. Oscillatory phytoplankton population dynamics are influenced by nutrient and light availability (Becker *et al*., 2018; Hu *et al*., 2018; Hunter-Cevera *et al*., 2020; C. Li *et al*., 2022; Malerba *et al*., 2021; Mattern *et al*., 2022; Ribalet *et al*., 2015; Tsakalakis *et al*., 2022; Welkie *et al*., 2018) and by density- and size-dependent feedback processes with other community components (Acevedo-Trejos *et al*., 2016; Chen & Smith, 2018; De Corte *et al*., 2019; Hunter-Cevera *et al*., 2014; Sosik *et al*., 2003; Talmy *et al*., 2019b; Taniguchi *et al*., 2014). As a result, these interactions lead to diel oscillations in related ecological processes, including grazing rates, viral infection rates, and viral activity (Arias *et al*., 2020; Aylward *et al*., 2017; Connell *et al*., 2020; Demory *et al*., 2020; Mruwat *et al*., 2021; Ng & H. Liu, 2016). The presence of diel oscillations often make it challenging to infer process from pattern, e.g., reduced population growth and/or increased mortality can have the same net effect on abundances (Hunter-Cevera *et al*., 2014; Mattern *et al*., 2022; Ribalet *et al*., 2015; Sosik *et al*., 2003).

Across oceanic basins, grazers and viruses are hypothesized to be the dominant drivers of phytoplankton loss (Mojica *et al*., 2016; Pasulka *et al*., 2015). However, estimating the relative contribution of viral-induced and grazing-induced mortality at a particular site remains challenging in the absence of additional ecosystem-specific process information (Beckett & Weitz, 2017, 2018; Calbet & Saiz, 2017; Mruwat *et al*., 2021; Talmy *et al*., 2019a). We focus our analysis on a near-surface Lagrangian parcel of water in the NPSG, sampled at high resolution at 15m depth over ten days in summer 2015 by the SCOPE HOE-Legacy 2A cruise (see Methods). The oligotrophic NPSG is numerically dominated by the unicellular cyanobacterium *Prochlorococcus*, the most abundant photosynthetic organism in the global oceans (Partensky *et al*., 1999). Prior work using the cellular iPolony method estimates that cyanophage, despite being highly abundant, contribute to <5% of total *Prochlorococcus* cellular losses per day (Mruwat *et al*., 2021). In parallel, analysis of food requirements to maintain heterotrophic nanoflagellate abundances suggest that grazing could account for the majority of daily *Prochlorococcus* cell losses (Connell *et al*., 2020). However, this quota method cannot rule out potentially significantly lower rates of grazing, especially if *Prochlorococcus* represent only a part of the diet of heterotrophic nanoflagellates. Grazing is expected to drive the flow of matter through marine food webs and out of the surface ocean ecosystem via export and subsequent sinking of fecal pellets (Azam *et al*., 1983). In contrast, viral infection and lysis is expected to shunt matter back into the microbial loop (Fuhrman, 1999; Weitz & Wilhelm, 2012; Wilhelm & Suttle, 1999), though significant lysis (e.g., during blooms) may lead to sticky aggregate production and increased export out of the surface ocean (Breitbart *et al*., 2018; Sullivan *et al*., 2017). Hence, disentangling the relative rates of viral-induced lysis and grazing can help inform estimates of the link between primary production and export.

Here, we use an ecological modeling and statistical fitting framework, combined with field observations, as a means to understand how observed *Prochlorococcus* dynamics are shaped by a combination of bottom-up and top-down forces in the NPSG. The multi-trophic models combine principles of virus-microbe interactions and grazing (Grossowicz *et al*., 2017; Mateus, 2017; Talmy *et al*., 2019a; Weitz *et al*., 2015; Wirtz, 2019) with light-driven forcing of cellular physiology(Ribalet *et al*., 2015; Sosik *et al*., 2003). Using Bayesian Markov chain Monte Carlo fitting methods, we compare *in silico* model dynamics with measured *in situ* ecological rhythms. We then use model-data fits across a range of ecological scenarios as a means to robustly estimate the contribution of viral-induced lysis and grazing to total *Prochlorococcus* mortality. As we show, model-data integration suggests the tight coupling of *Prochlorococcus* growth and loss over diel cycles in the NPSG is due primarily to the impact of grazing – and not viral-induced mortality. In doing so, we also find that additional loss factors beyond top-down control of *Prochlorococcus* may be ecologically relevant, raising new questions on governing mechanisms in surface ocean ecosystems.

## 2 Methods

### 2.1 Model overview

We developed a mechanistic mathematical model of an Ecological Community driven by Light including Infection of Phytoplankton (ECLIP). Our model includes dynamics of *Prochlorococcus*, grazers, and viruses, as well as *Prochlorococcus* division and loss, where the loss arises due to a combination of grazing, viral lysis, and other factors (see Figure 1a). In this model, viruses correspond to the abundances of T4- and T7-like cyanophages, known to primarily infect *Prochlorococcus*. Grazers represent heterotrophic nanoflagellates which feed on multiple prey types (Frias-Lopez *et al*., 2009), however the primary prey for heterotrophic nanoflagellates could be *Prochlorococcus* (Connell *et al*., 2020). We introduced flexibility in our framework to account for the extent to which *Prochlorococcus* constitutes the primary food source for the observed grazer class. To assess this uncertainty we investigated six grazer strategies, ranging from a “specialized” grazer class exclusively consuming *Prochlorococcus* cells (*γ* = 0 day^−1^) to models with increasing levels of generalism (*γ* = 0.01 to *γ* = 0.5 day^−1^) representative of grazers consuming additional prey, e.g., heterotrophic bacteria which are not explicitly integrated into the model. Mixotrophic nanoflagellates (Caron, 2017) were observed, but contribute less to the grazing pressure on the bacterial community compared to heterotrophic nanoflagellates (Connell *et al*., 2020). As it was not possible to differentiate abundance measurements of mixotrophic nanoflagellates from phototrophic nanoflagellates (Connell *et al*., 2020), we focus on grazing by heterotrophic nanoflagellates. Across the gradient of specialism-generalism grazer models, we searched for biologically feasible parameters using a model-data integration approach to generate dynamics consistent with observed population dynamics in the NPSG.

**Figure 1:**
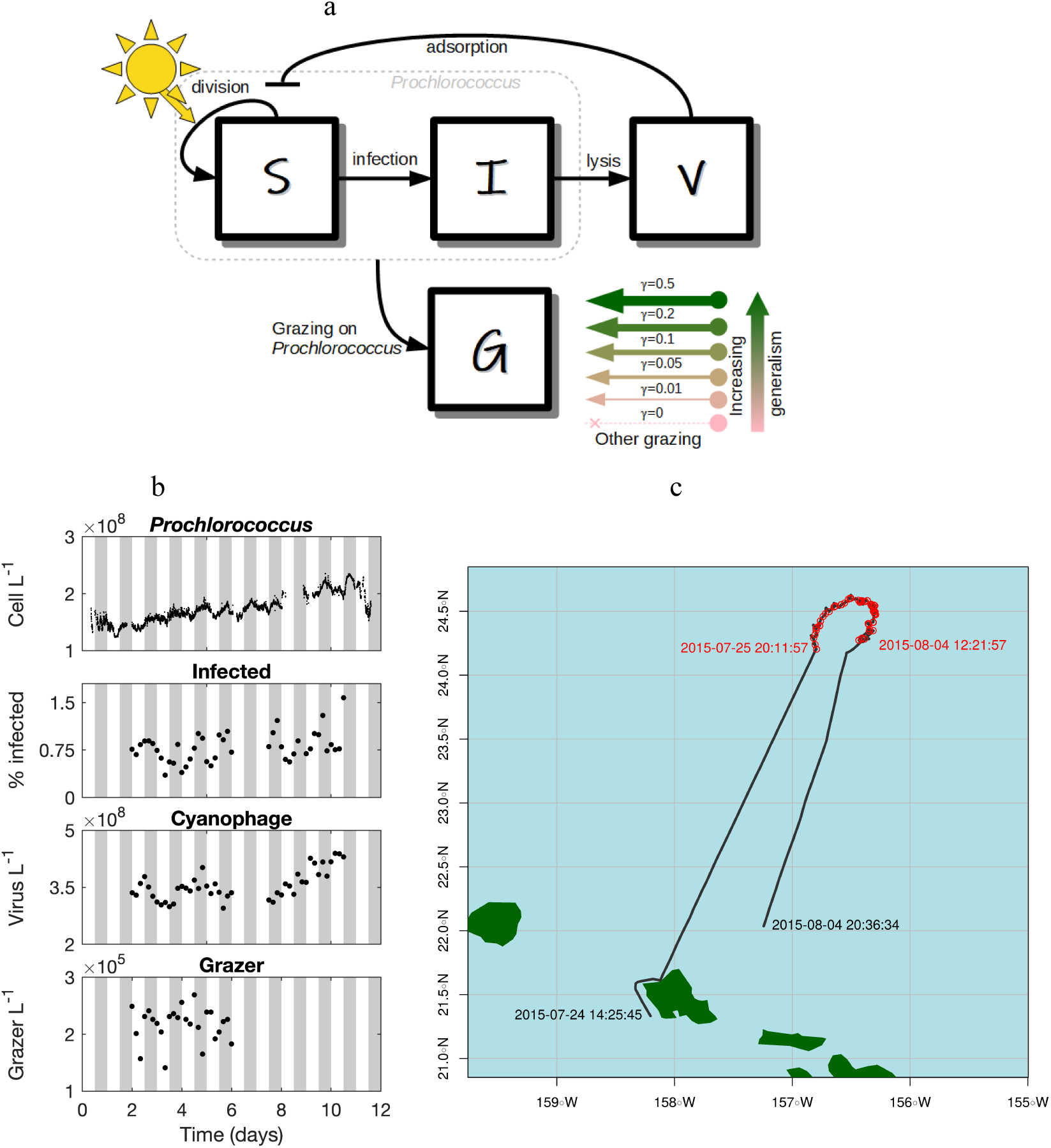
Community ecological model of viral and grazer mediated predation; and SCOPE HOE-Legacy 2A cruise field data. (a) *Prochlorococcus* are structured by infection status. Viruses (*V*) can infect susceptible *Prochlorococcus* cells (*S*) generating infected cells (*I*). Viral-induced lysis of infected cells releases virus particles back into the environment. Susceptible and infected *Prochlorococcus* cells are subject to grazing pressure from heterotrophic nanoflagellate grazers (*G*). Grazers may have a generalist strategy (e.g., grazing on heterotrophs, mixotrophs and phytoplankton not represented by *S* and *I*). We specify six models along this specialism-generalism gradient by setting a parameter *γ*. When *γ* = 0 heterotrophic nanoflagellate grazers act as specialists and only consume *Prochlorococcus*; and as *γ* increases, *Prochlorococcus* constitutes less of the diet of heterotrophic nanoflagellate grazers. Parameters and units are specified in Table S2. (b) Empirical population dynamics of *Prochlorococcus* cells, the percentage of *Prochlorococcus* cells infected with T4/T7-like cyanophage, the abundance of free-living T4/T7-like cyanophage, and the abundance of heterotrophic nanoflagellate grazers. (c) Cruise track and sampling stations. Local times (HST) for the start and end of recorded underway sampling (black line), and first and last sampling stations (red points) are annotated.

### 2.2 Ecological model of phytoplankton communities with viral and grazer mediated predation (ECLIP)

The ECLIP model represents *Prochlorococcus* cell division as a light-driven process where cell division is expected to occur at night (Binder & DuRand, 2002; Ribalet *et al*., 2015) and *Prochlorococcus* cell losses are controlled by viral lysis, grazing, and other density-dependent factors (Figure 1a). The *Prochlorococcus* population is divided into cells that are susceptible to viral infection (*S*) and cells that are infected (*I*) by viruses (*V*). Grazers (*G*) feed indiscriminately on both *S* and *I* classes. Abundance dynamics of *S*, *I*, *V* and *G* over time are described by the following equations:

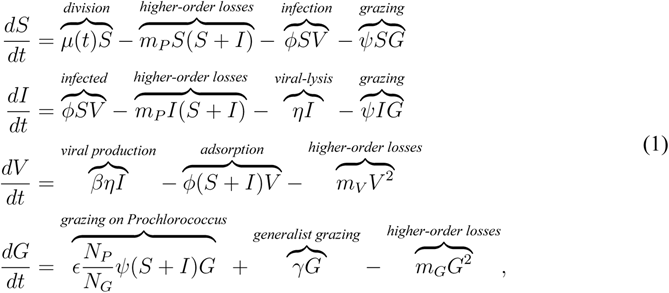

where

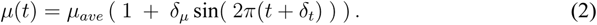

*Prochlorococcus* have a diel-driven division rate *µ*(*t*) whose proportional amplitude and phase are set by parameters *δ_µ_* and *δ_t_* respectively, and *t* = 0 represents 06:00:00 local time (*t* in days) (see Figure S1). *Prochlorococcus* have a nonlinear loss rate, *m_P_*, dependent on total *Prochlorococcus* abundance, implicitly representing niche competition (Berube *et al*., 2019). Viruses infect susceptible *Prochlorococcus* at rate *φ*. For each infection a burst size of *β* new virions are released into the environment upon cellular lysis following the latent period (average duration 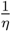). Grazing upon *Prochlorococcus*, at rate *ψ*, is non-preferential regarding infection status. Consumed *Prochlorococcus* biomass is converted into grazer biomass with Gross Growth Efficiency (GGE) *E* and assumed proportional to a nitrogen currency, given the nitrogen content in a *Prochlorococcus* cell (*N_P_*) and a grazer (*N_G_*), leading to an effective GGE of 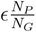. We introduce *γ* to denote the level of generalism in grazing, where *γ* represents net additional gains to the grazer from non-*Prochlorococcus* prey sources after accounting for respiratory costs. A specialist strategy has *γ* = 0 day^−1^, assuming that *Prochlorococcus* cells are the only grazer prey source. In contrast, we represent five generalist strategies via *γ* = 0.01 (very low), 0.05 (low), 0.1 (medium), 0.2 (high), or 0.5 (very high) day*^−^*^1^, implying that grazers have a net positive growth rate even in the absence of *S* or *I* via consumption of other phytoplankton, heterotrophic bacteria, or grazers (intraguild predation). The degree that grazers act as generalists, rather than specialists on *Prochlorococcus*, depends on other life-history traits (equation S20). Grazer and viral losses are characterized by nonlinear loss terms (with rates *m_G_* and *m_V_*) to avoid structurally biasing the model to favor one *Prochlorococcus* predator type (Talmy *et al*., 2019a) and to avoid competitive exclusion between grazers and viruses. Linear loss terms were excluded to reduce the number of free parameters to estimate via inference. See further details in the Supplementary Information (see Table S2 for parameter definitions and Table S3 for specification of parameter priors).

### 2.3 *in situ* measurements

We use data collected from the Summer 2015 SCOPE HOE-Legacy 2A cruise (Figure 1b-c). Measurements of total *Prochlorococcus* abundance were made every *≈*3 minutes, with measurements of infected cells (infected by either T4- or T7-like cyanophages), total virus particles of either T4- or T7-like cyanophages, and heterotrophic nanoflagellate grazers collected at 4 hr intervals over a multi-day period aboard the R/V Kilo Moana. In all figures, the 12 days shown represent 06:00:00 24th July 2015 to 06:00:00 August 5th 2015 local time.

#### Prochlorococcus cell abundance

SeaFlow – a shipboard *in situ* flow cytometer – continuously measures forward scattering, red and orange fluorescence intensities of particles ranging from ~0.4 to 4 *µ*m in diameter from underway samples (continuously pumped surface seawater from ~7 m depth) every 3 minutes. A combination of manual gating and statistical methods was used to identify *Prochlorococcus* based on forward scatter (457/50 bandpass filter), orange fluorescence (572/28 bandpass filter) and red fluorescence (692/40 band-pass filter) relative to 1-*µ*m calibration beads (Invitrogen F8823). Individual cell diameters were estimated from SeaFlow-based light scatter by applying Mie light scatter theory to a simplified optical model, using an refractive index of 1.38 (Freitas *et al*., 2020; Ribalet *et al*., 2019a,b). Data obtained via Simons CMAP (Ashkezari *et al*., 2021).

#### Virus abundance and infection

Samples for virus abundance and infection were collected every 4 hours at 15 m depth using a CTD-rosette equipped with 12 L niskin bottles (Mruwat *et al*., 2021). Samples for cyanophage abundances (40 mL) were filtered through a 0.2 *µ*m syringe top filter, flash frozen, and stored at −80°C. Samples for infected cells (40 mL) were filtered through a 20 *µ*m nylon mesh, fixed with electron microscopy grade glutaraldehyde (0.125% final concentration), incubated for 30 minutes in the dark at 4°C, flash frozen, and stored at −80°C. Cyanophage concentrations were analyzed using the polony method for T7-like (Baran *et al*., 2018) or T4-like (Goldin *et al*., 2020) cyanophage families. Virally infected *Prochlorococcus* was quantified using the iPolony method (Mruwat *et al*., 2021) in which *Prochlorococcus* cells were sorted with a BD Influx cytometer and screened for intracellular T4-like and T7-like cyanophage DNA.

#### Heterotrophic nanoflagellates

Samples for nanoplankton (protists 2-20 *µ*m in diameter) abundances were collected every 4 hours at 15 m depth (Connell *et al*., 2020). Subsamples were preserved with formaldehyde (final concentration 1%, final volume 100 mL) and stored at 4°C. Slides were prepared from preserved samples within 12 hours of sampling by filtering 100 mL subsamples down to ~1 mL onto blackened 2 *µ*m, 25 mm polycarbonate filters and staining the samples with 50 *µ*L of a 4’-6’diamidino-2-pheylindole (DAPI, Sigma-Aldrich, St. Louis, MO) working solution (1 mg mL*^−^*^1^) for 5-10 minutes in the dark (Sherr *et al*., 1993). Stained samples were filtered and rinsed; filters were placed on glass slides with a drop of immersion oil and coverslip, then sealed with clear nail polish. Slides were stored at −20°C. Heterotrophic nanoplankton abundances were counted using epifluorescence microscopy from triplicate slides, and differentiated from photo/mixotrophic nanoplankton by the lack of chlorophyll *a* autofluorescence in plastidic structures when viewed under blue-light excitation (Connell *et al*., 2020).

### 2.4 Model-data integration

To constrain ECLIP models to data, we used Markov Chain Monte Carlo (MCMC), implemented in the Turing package (Ge *et al*., 2018) in Julia (Bezanson *et al*., 2017). MCMC is a class of Bayesian inference algorithms that aims to infer model parameter probability distributions given the structure of model equations, data, and prior parameter distributions. We used the No-U-Turn Sampler to sample posterior distributions (Hoffman, Gelman, *et al*., 2014). Further details are in the Supplementary Information.

## 3 Results

### 3.1 Fitting ECLIP to field measurements of *Prochlorococcus*, cyanophages, and grazers

Timeseries data from the SCOPE HOE-Legacy 2A cruise (see Figure 1b) reveals *Prochlorococcus* abundances are periodic, peaking at night and reaching their minima during the day (Mruwat *et al*., 2021). Population abundances of heterotrophic nanoflagellates fluctuate with unclear periodicity (Connell *et al*., 2020), as do the abundances of T4- and T7-like cyanophage (Mruwat *et al*., 2021). In contrast, the fraction of cells infected by T4- and T7-like viruses (Mruwat *et al*., 2021) are periodic, peaking at night. We recapitulated these periodicity analyses in Supporting Information Section S1. This periodicity suggests the potential for diel-driven emergent synchronization in the food web, similar to community-wide metabolism in the NPSG (Muratore *et al*., 2022).

To explore potential coexistence dynamics of *Prochlorococcus*, viruses and grazers, we fit the ECLIP model via MCMC given biologically realistic parameter bounds (see Table S3 for priors, Figure S2 for division-associated priors, and the Supplementary Information for MCMC fitting details). The models are fitted against detrended empirical data, so for visualization we add this trend to the model simulations. The fitting of ECLIP with differing levels of grazer generalism are shown in Figure 2. All ECLIP models were able to simultaneously reproduce the magnitudes of the different timeseries, producing fits with similar log-likelihoods (Figure S9) while exhibiting statistical evidence of convergence (Figure S5 and Figure S8), even if it is not feasible to identify a particular, preferred level of grazer generalism. In sum, a range of nonlinear mathematical models including feedback between cyanobacteria, cyanophage, and grazers can jointly recapitulate multi-trophic population dynamics in the NPSG. Despite fitting overall magnitudes and oscillations in *Prochlorococcus* abundances, ECLIP underestimated the strength of oscillations in infected cells (an issue we return to later in the Results).

**Figure 2:**
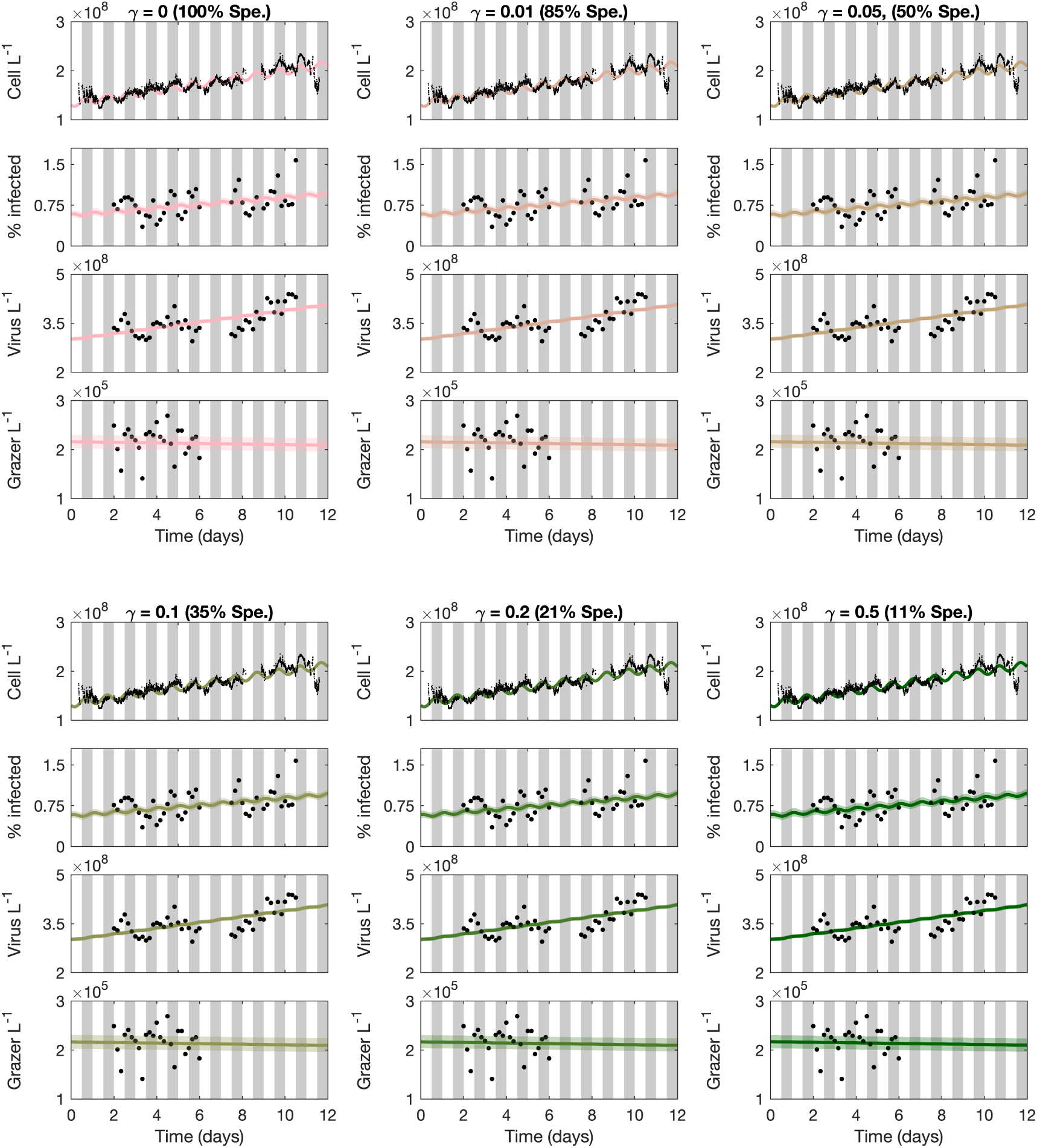
Models across the specialism-generalism gradient fit empirical data. ECLIP models (lines) are compared against empirical data (points). Model lines represent the median MCMC solution within 95% CI range found by the converged chains, shown as bands with colours representing the choice of *γ*. Data signals include *Prochlorococcus* cell abundances (top), the percentage of infected *Prochlorococcus* cells, the abundance of free viruses and the abundance of heterotrophic nanoflagellate grazers (bottom). The models were fitted against detrended data; for visualization we have added these trends to the model solutions. Grey bars indicate nighttime. Model solutions with: (a) *γ* = 0 (grazers act as specialists), (b) *γ* = 0.01, (c) *γ* = 0.05, (d) *γ* = 0.1, (e) *γ* = 0.2, (f) *γ* = 0.5 day*^−^*^1^. The degree of grazer specialism (Spe.) is shown in parentheses above each subplot.

### 3.2 Interpretation of ecological mechanisms underlying model-data fits

The equivalence in model-data fits across a spectrum of grazer generalism suggests that differentiating model mechanisms requires inspection of posterior parameter fits. Posterior parameter values are shown in Figure 3, with full details on division function fits in Figure S3 and comparison with priors in Figure S4. Most life-history traits converge to similar parameter regimes across ECLIP models, with a notable exception: a systematic trend in the grazer loss parameter *m_G_*, reflecting a trade-off between grazer losses and gains via increasing *γ*. The inferred grazer loss rates correspond to grazer residence times of between 16.81 (95% CI: 15.37-20.19) days in the specialist model to 1.82 (95% CI: 1.8-1.87) days in the most generalist model when *γ* = 0.5 per day. These residence times are consistent with the range of estimated heterotrophic nanoflagellate doubling times in the Mediterranean Sea of 4–20 days (Christaki *et al*., 2001). Corresponding virus residence times were estimated as 1.16 days (95% CI: 0.59-5.3 days) in the specialist model and 1.17 days (95% CI: 0.57-8.32 days) when *γ* = 0.5 (see Figure S10). MCMC posterior distributions appear tight (e.g., for *µ_ave_*, *δ_t_*, and *φ*) or loose (e.g., for *β*, *m_P_* and *m_V_*) suggesting differing parameter space sensitivities (Gutenkunst *et al*., 2007). Figure 4 shows that *γ* corresponds to a grazer specialism-generalism gradient in the inferred ECLIP models, with the relative contributions of *Prochlorococcus* consumption to grazer growth rates decreasing with increasing generalism, as expected (note life-history trait interdependence, as in equation S20, did not guarantee this result). However, absolute per capita grazer consumption of *Prochlorococcus* only varied modestly (*≈*0.04-0.075 day*^−^*^1^) between models. Hence, we interpret these findings to mean that per-capita grazing mortality of *Prochlorococcus* is relatively invariant to model choice and can be inferred robustly from model-data fits.

**Figure 3:**
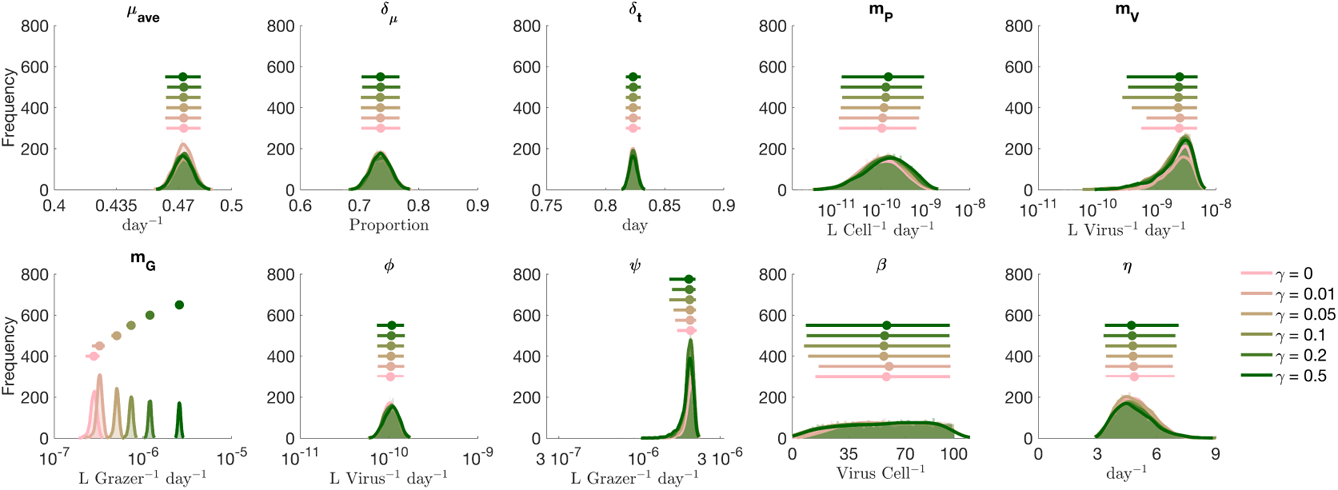
Differences in inferred life-history traits across the specialist-generalist gradient. (a-j) Parameter posterior distributions for different ECLIP models. Parameters are (a) *µ_ave_*: average *Prochlorococcus* division rate, (b) *δ_µ_*: division rate amplitude, (c) *δ_t_*: phase of division rate, (d) *m_P_* : higher order *Prochlorococcus* loss rate, (e) *m_G_*: higher order viral loss rate, (f) *m_G_*: higher order grazer loss rate, (g) *φ*: viral adsorption rate, (h) *ψ*: grazer clearance rate, (i) *β*: viral burst size, and (j) *η*: viral-induced lysis rate. Jittered median (dot) and 95% CI range (horizontal line) for each of the models are shown above density plots. Full details of parameter bounds are shown in Table S2; see Supplementary Information for more details.

**Figure 4:**
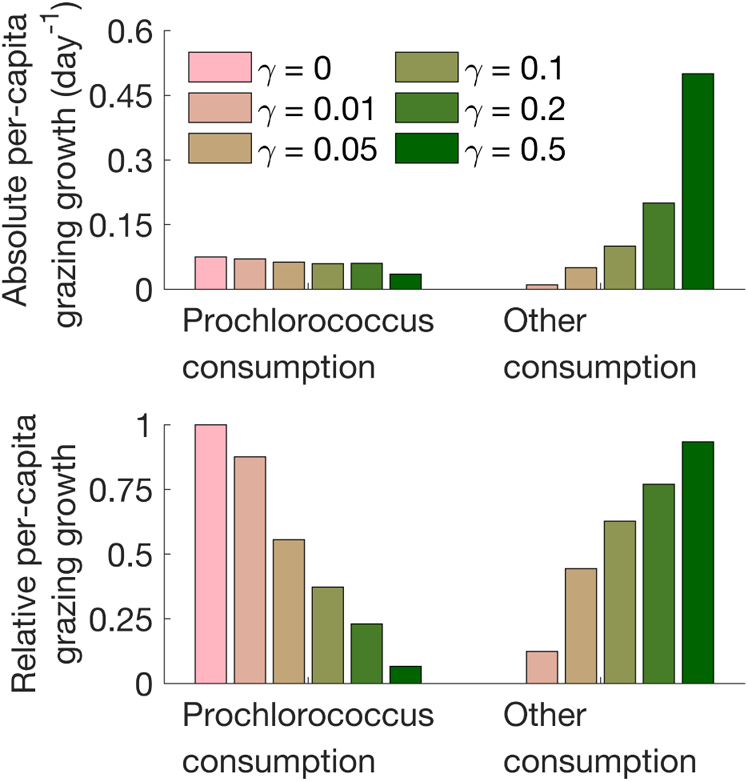
Model differences across the specialist-generalist gradient. (a-b) Inferred grazer growth attributable to consumption of *Prochlorococcus* or other sources (see equation S20) across models.

### 3.3 Partitioning *Prochlorococcus* losses between top-down and other effects

We analyzed the predicted partitioning of *Prochlorococcus* mortality among grazing by heterotrophic nanoflagellates, viral-induced lysis by T4- and T7-like cyanophages, and other sources of mortality, using the inferred ECLIP models. We used posterior estimates from model-data fits to estimate total *Prochlorococcus* loss rates:

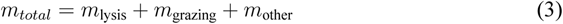

where: *m*_lysis_ = *ηI*, *m*_grazing_ = *ψ*(*S* +*I*)*G*, *m*_other_ = *m_P_* (*S* +*I*)^2^ (see Supplementary Information for details). The proportion of each mortality process is calculated as the average ratio of the component mortality rate relative to that of the total over the empirical time-series. Mortality partitioning suggests 87-89% (with extents of 95% confidence intervals ranging 66-96%) of *Prochlorcoccus* losses were ascribed to grazing, 6% (with extents of 95% confidence intervals ranging 4-9%) to viral lysis and 4.5-6.6% (with extents of 95% confidence intervals ranging from less than 1% to 27%) to other mechanisms. The distribution of losses by category are shown in Figure 5, with the top six rows in each panel denoting distinct grazer generalism levels, from *γ* = 0 to *γ* = 0.5. Inferred mortality estimates from ECLIP were relatively invariant regardless of the grazer generalism level. Hence, our estimates of mortality partitioning are robust to model choice, including the finding that grazer-induced losses predominate when jointly estimating the collective effects of grazers and viruses on multitrophic population dynamics. Notably, other forms of loss may be as ecologically relevant as viral-induced mortality to daily *Prochlorococcus* losses.

**Figure 5:**
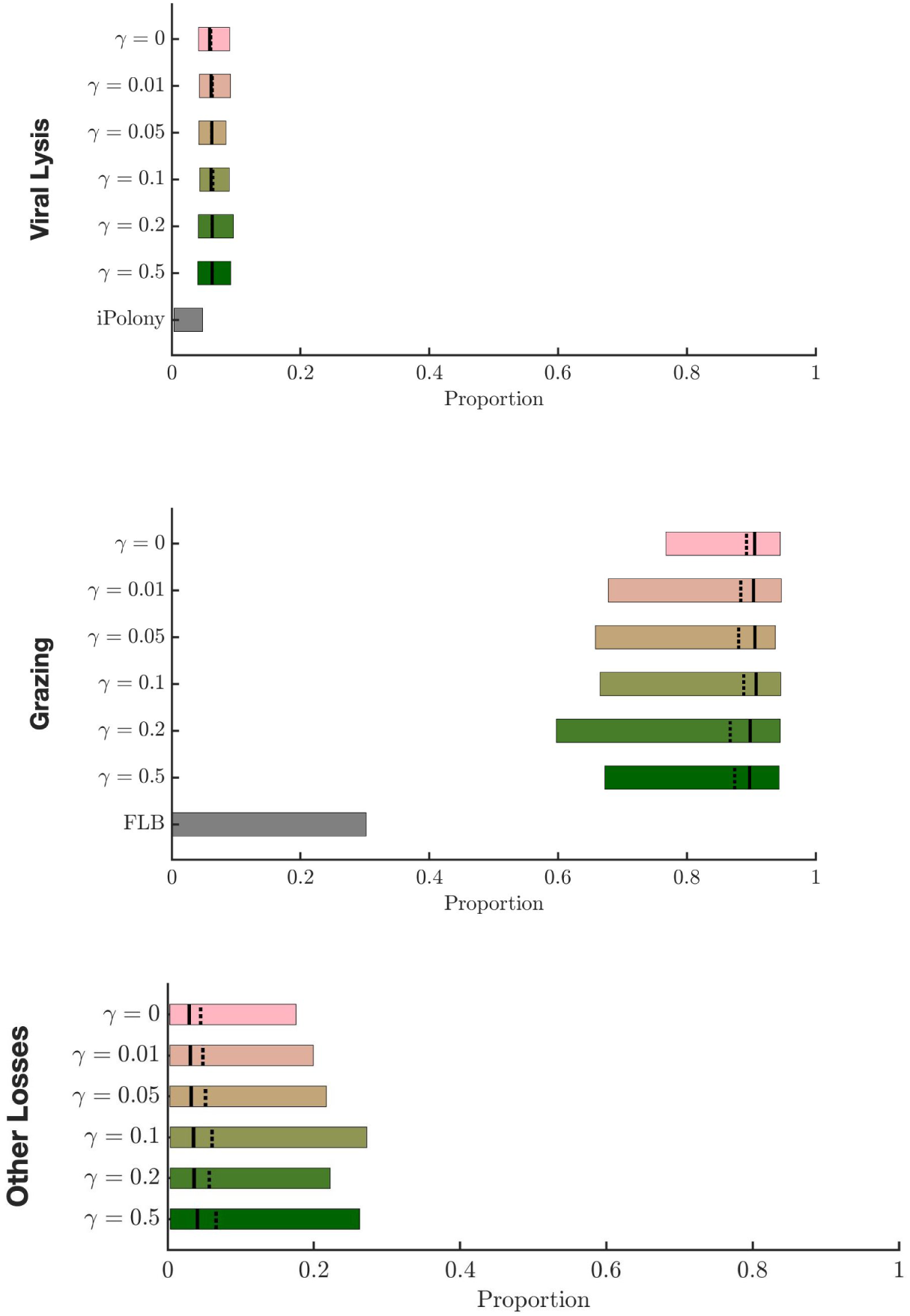
Relative importance of viral lysis, grazing, and other effects on total *Prochlorococcus* mortality. The proportion of mortality partitioned between viral-induced lysis (top panel), grazing (middle panel), and other (bottom panel) sources for the ECLIP models and other measures of relative mortality. For ECLIP the results from all chains are shown. Bars in these panels denote mortality rate proportions associated with the 95% confidence intervals, where the mean and median are shown by solid and dashed lines, respectively. Other plotted measures of relative mortality are given via direct measurements of viral infection (iPolony), and Fluorescently Labelled Bacteria (FLB) incubation measurements (see Supplementary Information Section S2 for details).

### 3.4 Contrasting *Prochlorococcus* loss estimates

ECLIP model-data integration simultaneously infers the putative daily loss of *Prochlorococcus* due to grazing, viral lysis and other loss mechanisms. These joint estimates can be compared to alternative methods that estimate viral infection or grazing losses, albeit one factor at a time. Multiple approaches exist to infer *Prochlorococcus* loss rates *in situ*. For viral lysis, we consider two methods: (i) encounter theory; (ii) iPolony estimates. Conventional ‘encounter’ estimators use biophysical theory to estimate an upper-limit of size-dependent contact rates. However, encounter need not imply a successful adsorption and lysis event, hence the realized level of lysis is often significantly less than expected from biophysical limits (Talmy *et al*., 2019b). In contrast, the iPolony method quantifies the fraction of host cells infected by a target phage, which can be combined with estimates of viral latent periods and cell division rates to infer loss rates (Mruwat *et al*., 2021). Regarding grazing, we consider three methods: (i) encounter theory; (ii) quota-based theory; (iii) flourescently-labeled bacteria (FLB) estimates. For grazers, size-dependent encounter rate theory and quota-based theories use biophysical contact rates and allometrically derived elemental growth requirements, respectively, to estimate grazing-induced loss rates. Theoretical estimates via encounter and quota methods have significant variability, in part due to life-history trait uncertainties. FLB is a direct method, insofar as it quantifies surrogate prey uptake (*Dokdonia donghaensis*, see Connell *et al*., 2020) as a proxy for cyanobacteria uptake rates.

Figure 5 compares *joint* ECLIP-inferred relative mortality estimates with *one-factor* estimates from field-based iPolony measurements and fluorescently labelled bacterial (FLB) uptake estimates (for details of these and alternative, theoretical methods (encounter and quota), see Supplementary Information S2). For viral lysis, one-factor estimates using encounter theory do not constrain daily loss rates, given that contact-limited lysis rates are compatible with nearly 100% of observed *Prochlorococcus* loss; but decreases in contact rates, efficiency of adsorption, and inefficiency of infection post-lysis lead to poorly constrained lower limits. In contrast, quantitative estimates of infection processes in the NPSG via the iPolony method suggest viral-induced lysis by T4- and T7-like cyanophages contribute a comparatively small amount to *Prochlorococcus* cell losses (<5%). Likewise, accounting for observed grazer abundances and biophysically plausible grazing rates suggests grazing could explain daily *Prochlorococcus* losses alone. But, lower limits of theory based estimates of grazing-induced mortality – as was the case for viral-induced mortality – are poorly constrained given uncertainties in grazing efficiency. For example, empirically-derived FLB estimates suggest grazing by heterotrophic nanoflagellates accounts for *~*30% of total *Prochlorococcus* loss rates (although we note the original FLB experiment was designed to differentiate relative grazing by mixotrophic and heterotrophic nanoflagellates). If the iPolony and FLB methods were unbiased, then the majority of loss rates would be unaccounted for by top-down effects.

Instead, our mortality estimate comparison constrain the magnitude of distinct loss factors. Like the iPolony method, our joint ECLIP-inferred estimates suggest that viral-induced lysis is responsible for a small proportion of *Prochlorococcus* losses in the NPSG. We note that ECLIP-inferred viral lysis estimates are low but are *~* 3% higher than those inferred from iPolony (see Discussion). In contrast, joint ECLIP-inferred estimates suggest grazing by heterotrophic nanoflagellates represents the majority of *Prochlorococcus* cell losses across ECLIP – far above that inferred via FLB experiments. Notably, estimates of *Prochlorococcus* losses via grazing were robust to changes in grazer specialism, further suggesting FLB-derived estimates under-represent *in situ* grazing (which could represent biological or methodological uncertainty as hypothesized in Connell *et al*., 2020). Additionally, the model-inferred combination of viral-induced lysis by T4- and T7-like cyanophages and grazing by heterotrophic nanoflagellates typically does not sum to equal 100% of total *Prochlorococcus* daily cell losses. Instead, model-data integration suggests other sources of *Prochlorococcus* cell loss account for *≈*6% of daily losses (with 95% confidence intervals ranging from less than 1% to 27%). Together, both model-data fits and independent estimates of top-down mortality suggest other loss processes beyond grazing by heterotrophic nanoflagellates and lysis by T4/T7-like cyanophages may be critical in shaping daily phytoplankton rhythms.

### 3.5 Capturing diel periodicity of infected *Prochlorococcus*

The multi-trophic ECLIP model incorporating light-driven photosynthesis resolved the diel periodicity of total cell counts of *Prochlorococcus* across a gradient of grazer consumption strategies (see Figure 2). However, the ECLIP model did not recapitulate the magnitude of the observed periodicity of infected cells (as shown in Supporting Information S1). One potential reason for this gap is that we did not incorporate the potential for plasticity in viral traits into ECLIP, in contrast to previous work which shows that cyanophage exhibit light-dependent adsorption rates to *Prochlorococcus* (Demory *et al*., 2020; R. Liu *et al*., 2019). Diel-dependent adsorption may reflect changes in both cell physiology and cell size as *Prochlorococcus* cells grow during the day in G1 phase before synthesising DNA in S phase and transitioning to G2 phase in preparation to divide at night – larger cells are expected to have larger rates of adsorption (Talmy *et al*., 2019b) and darkness can modulate and arrest transitions through cell cycle phases (Hynes *et al*., 2015; Vaulot, 1995; Vaulot & Marie, 1999) which in turn could modulate viral infection (Ni & Zeng, 2016). Hence, we modified the core ECLIP model to include a time-dependent step-wise adsorption rate such that adsorption is 50% lower at dawn (midnight to noon) and 50% higher at dusk (noon to midnight) than the initially inferred adsorption rates (preserving the mean adsorption rates used in Figure 2). The emergent community dynamics preserve the timing and magnitude of oscillations in *Prochlorococcus* populations while also inducing oscillations in infected cells (see Figure 6 for the case with *γ* = 0.05 and Figure S11 for the full suite of grazer strategies). Hence, we find that it is possible to recapitulate the daily community time series insofar as we incorporate both light-dependent cellular and viral traits.

**Figure 6:**
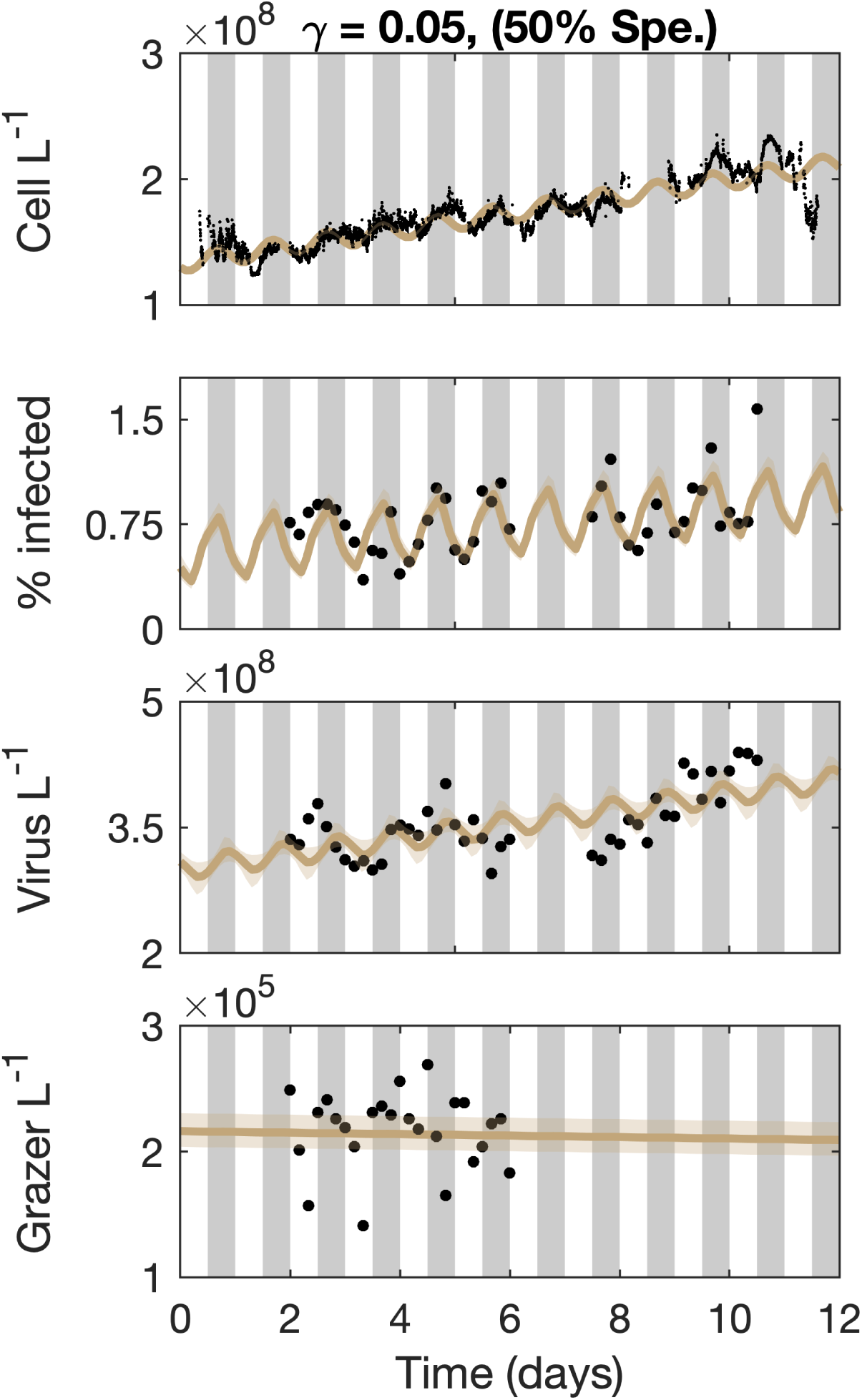
Diel-dependent adsorption rates improve fits to infected cells. ECLIP model solutions with *γ* = 0.05 and diel-dependent adsorption rates are compared against empirical data in black. Model lines represent the median MCMC solution within 95% CI range found by the converged chains, shown as bands. Data signals include *Prochlorococcus* cell abundances (top), the percentage of infected *Prochlorococcus* cells, the abundance of free viruses and the abundance of heterotrophic nanoflagellate grazers (bottom). The models were fitted against detrended data; for visualization we have added these trends to the model solutions. Grey bars indicate nighttime. The degree of grazer specialism (Spe.) is shown in parentheses above the plot.

## 4 Discussion

We developed and analyzed a multitrophic community ecology model (ECLIP) consisting of *Prochlorococcus*, viruses, and grazers to investigate feedback mechanisms and ecological drivers of oligotrophic ocean microbial population dynamics on diel timescales. ECLIP can recapitulate the dynamical coexistence of cyanobacteria, viruses infecting cyanobacteria, and grazers population abundances in the NPSG. By combining model-data fits with direct measurements of mechanistic interactions we infer that grazing rather than viral-induced lysis predominates in shaping *Prochlorococcus* mortality in NPSG surface waters. We also find that the combination of lysis and grazing does not fully account for daily *Prochlorococcus* losses. Instead, model-inference suggests a non-trivial loss rate due to mechanisms beyond that of top-down lysis and grazing.

Overall, model-data fitting to NPSG measurements enabled us to examine how much *Prochlorococcus* mortality can be ascribed to viral lysis, heterotrophic nanoflagellate grazing, or other loss processes. In partitioning *Prochlorococcus* mortality, we found different outcomes across model scenarios and independent auxiliary estimates (Figure 5). Indirect estimates via encounter or quota-based theory are poorly constrained and limited by our current knowledge of ecological life-history traits. However, fitting ECLIP to field data resulted in more constrained mortality estimates. Viral-induced mortality of *Prochlorococcus* in this system is relatively weak, consistent with prior estimates (Mruwat *et al*., 2021). Low levels of viral-induced *Prochlorococcus* (and *Synechococcus*) mortality have also been found in the Sargasso Sea (Matteson *et al*., 2013), and the Mediterranean and Red Sea (Mruwat *et al*., 2021). This could be a defining characteristic of viral impacts on cyanobacteria in oligotrophic gyres – as opposed to more dynamic ocean regions where viral mortality can be considerably more substantial (Carlson *et al*., 2022; Fuhrman & Noble, 1995).

Direct mortality estimates from grazing incubation experiments and infected cell measurements also provide evidence that heterotrophic nanoflagellate grazing and T4- and T7-like viral-induced mortality do not account for all *Prochlorococcus* losses in the NPSG. Quantifying the relative importance of mortality processes beyond conventional top-down effects (grazing and lysis) is critical for understanding how grazers and viruses contribute to mortality and energy transfer in marine microbial communities (Mojica *et al*., 2016; Pasulka *et al*., 2015; Ribalet *et al*., 2015; Talmy *et al*., 2019a). Interestingly, our analysis suggested lower levels of grazing mortality using FLB measurements compared to those inferred via ECLIP. This suggests grazers do not uptake this biological tracer at the same rate as *Prochlorococcus* – potentially reflecting differences in chemical composition, size, or experimental conditions (Connell *et al*., 2020).

The finding that model-data fits impute other sources of mortality as quantitatively significant suggests other feedback mechanisms should be included in model representations of marine surface community dynamics. A missing component in our modeling framework is the effects of mixotrophic nanoflagellates (Connell *et al*., 2020; Q. Li *et al*., 2021, 2022; Sanders & Porter, 1988; Stoecker *et al*., 2017) which are likely the main source of additional losses. We did not include mixotrophs in our framework due to experimental difficulties in separating phototrophic from mixotrophic nanoflagellates, and theoretical challenges of appropriate physiological modeling. However, surface ocean phytoplankton losses plausibly include factors beyond grazing and viral-induced lysis (Aguilera *et al*., 2021; Brum *et al*., 2014). In Supplementary Information S4 we review potential mechanisms contributing to the unaccounted losses of *Prochlorococcus*, beyond those from heterotrophic nanoflagellate grazing and T4- and T7-like viral-induced lysis. These include ecological feedbacks leading to distinct functional and/or light-driven responses, aggregation and/or sinking, stress, population heterogeneity, and the possibility of having missed other top-down mortality. Similarly, our measurements may miss population heterogeneities within *Prochlorococcus* masking our ability to interpret average per-capita mortality. Investigating alternative mechanisms of *Prochlorococcus* losses may improve understanding of how biomass and nutrients are transferred through marine food webs. Further investigation and characterization of growth dynamics may also be warranted, as mischaracterization may impact our ability to infer mortality processes – note, at depth *Prochloroccocus* has recently been shown to rely on mixotrophic strategies (Wu *et al*., 2022).

The ECLIP framework comes with caveats, despite inclusion of multiple populations and interactions. First, we focused on the impacts of direct, light-driven forcing of cyanobacterial division – hence, oscillations arising in other components reflect a combination of instabilities that can arise in nonlinear population models as well as the cascading impacts of such oscillations on the community. Unlike other picoplankton modeling efforts including generic loss terms, e.g., Hunter-Cevera *et al*., 2014; Hynes *et al*., 2015; Mattern *et al*., 2022; Ribalet *et al*., 2015, ECLIP can infer loss partitioning between grazing, lysis, and other processes. However, ECLIP does not explicitly capture size-structured processes which are important drivers of growth (Hunter-Cevera *et al*., 2014; Hynes *et al*., 2015; Mattern *et al*., 2022; Ribalet *et al*., 2015) and other ecological interactions (Talmy *et al*., 2019b). Additionally, light-driven forcing of division does not fully account for variability in processes such as nutrient content (Lopez *et al*., 2016; Muratore *et al*., 2022; Vislova *et al*., 2019), and metabolic state (Muratore *et al*., 2022). While these attributes are not specifically modeled, they may have bearing on inferring life-history traits. ECLIP provides a complementary framework for understanding marine microbial ecology; and we hope future efforts will attempt to blend these types of models e.g., (Beckett *et al*., 2021). Direct incorporation of diel impacts on grazing (Arias *et al*., 2020; Connell *et al*., 2020; Groussman *et al*., 2021; Ng & H. Liu, 2016) or viral traits (e.g., adsorption) (Demory *et al*., 2020; Thamatrakoln *et al*., 2018) may be required to mechanistically understand population dynamics on sub-daily timescales – and particularly the magnitude of oscillations, including infected cell abundances. Second, we have used two focal processes to examine how carbon and other nutrients in basal picoplankton are transferred, either up the food chain via grazing, or retained in the microbial loop via viral lysis (aka the viral shunt) (Fuhrman, 1999; Wilhelm & Suttle, 1999). This dichotomy reflects potential tension regarding the extent to which primary production stimulates the biological pump requiring further investigation. Model extensions could include mechanistic process of export explicitly, whether through coupling grazing to the generation of particles and/or examining the extent to which viral lysis generates aggregates which can be exported to the deep oceans via the viral shuttle (Sullivan *et al*., 2017; Weinbauer, 2004). Finally, our work has identified a potential accounting challenge in quantifying the balance of *Prochlorococcus* growth and losses. Despite the daily growth and division of cells, overall abundances remain tightly constrained – our work suggests this constraint depends on factors beyond loss ascribed to T4- and T7-like cyanophage and nanoflagellate grazers.

In summary, the ECLIP multitrophic modeling framework provides opportunities to disentangle putative mechanisms underlying the control of microbial surface ocean populations. The model provides support for the dominant role of grazers in controlling *Prochlorococcus* in the NPSG, that relatively high viral abundances can be compatible with relatively low infection (and mortality) rates, while also identifying a key direction for future work: identifying the potentially ‘missing mortality’ at the base of the marine food web. Moving forward, *in situ* observations are needed to probe aggregation and sinking, autolysis, programmed cell death, or other forms of loss of *Prochlorococcus* and to understand the feedbacks of coupled variation in cyanobacterial growth and loss in a changing ocean.

## Acknowledgements

We thank the captain and crew of the R/V Kilo Moana for their effort and assistance on the Hawaii Ocean Experiment Legacy II cruise (KM1513); we thank Michael Follows for feedback that helped improve the manuscript, and we thank Jeremy Harris for code review. This work was supported by grants from the Simons Foundation (no. 549894 to J.R.C., no. 721231 to J.S.W, and no. 329108 to E.V.A., D.A.C., D.L., A.E.W. and J.S.W). This is a contribution of the Simons Collaboration on Ocean Processes and Ecology (SCOPE). This research was supported in part through research cyberinfrastructure resources and services provided by the Partnership for an Advanced Computing Environment (PACE) at the Georgia Institute of Technology, Atlanta, Georgia, USA.

## Data availability

Code for ECLIP and performing Bayesian parameter inference was written and run in Julia, with plotting performed in MATLAB. Core code for running model simulations, analysis and plotting is archived on Zenodo: https://doi.org/10.5281/zenodo.7937657 (Beckett *et al*., 2023).

## Supplementary Information

### S1 Timeseries diel analysis

We used the R package RAIN (Thaben & Westermark, 2014) (version 1.34.0) to evaluate whether the empirical *in situ* timeseries exhibited diel periodicity. RAIN uses non-parametric methods to evaluate periodicity and is optimized to focus on evenly sampled datasets with fewer than 100 measurements. In all cases we first detrended the timeseries. For *Prochlorococcus* cells, the size of data goes well above what is computationally feasible within RAIN, and sample times are not evenly spaced. Instead, we used the Lomb-Scargle Periodogram (Ruf, 1999) implemented in R package *lomb* (version 2.1.0) to estimate the period (found to be between 0.977 and 1.023 days). Table S1 shows the results from the RAIN and Lomb-Scargle analyses. The timeseries of *Prochlorococcus* and infected *Prochlorococcus* was statistically significantly diel (at a level of p<0.05), the timeseries for T4- and T7-like viruses, and heterotrophic nanoflagellates was not statistically significantly diel (at a level of p>0.05).

**Table S1:** Diel timeseries analysis using Lomb-Scargle Periodogram for *Prochlorococcus* and RAIN for other timeseries, testing for the probability of whether timeseries are expected to exhibit periodicity of 1 day by chance.

| Timeseries | Method | *p*-value | Significantly diel? |
| --- | --- | --- | --- |
| *Prochlorococcus* | Lomb-Scargle Periodogram | 3.056296e-232 | ✓ |
| Infected cells | RAIN | 0.01042877 | ✓ |
| % infected cells | RAIN | 0.002237049 | ✓ |
| Virus | RAIN | 0.9064921 | × |
| Grazer | RAIN | 0.1904631 | × |

### S2 Alternative mortality estimates

#### S2.1 Modeling encounter rates and allometry

##### S2.1.1 Viral encounter rate

Both *Prochlorococcus* and their viruses diffuse and the rates at which they may encounter each other can be estimated using biophysical models (Berg & Purcell, 1977; Murray & Jackson, 1992; Talmy *et al*., 2019b). We assume the maximum encounter rate between diffusing spherical particles of two different sizes, *r_virus_* and *r_cell_* is given by the Smoulouski equation (Von Smoluchowski, 1917). However, an encounter does not always lead to adsorption (Murata *et al*., 2017; Talmy *et al*., 2019b) so we subject the maximum encounter rate to an efficiency of encounter term, *E_φ_*, as:

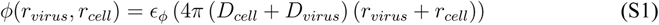

where diffusion of each spherical particle of radius *r* is given by the classical relation(Einstein, 1905):

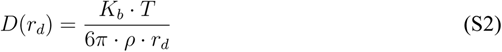

with *K_b_*, the Boltzmann constant, *T* is temperature (in Kelvin) and *ρ* is the dynamic viscosity of the medium. We assume *T* = 25°C (Karl & Church, 2014) and *ρ* = 9.96 *×* 10*^−^*^10^ kg *µ*m*^−^*^1^ s*^−^*^1^ (Talmy *et al*., 2019b). We assume that free viruses can adsorb to both susceptible and infected *Prochlorococcus* cells. Both cases result in losses of free virus particles. Susceptible cells that viruses adsorb to become infected. However, we assume that viruses that adsorb to infected cells do not alter the physiological responses of infected cells. As radius scales as *≈* [volume]^1/3^, both diffusion and encounter rate are size-dependent (Talmy *et al*., 2019b).

##### S2.1.2 Grazer encounter rate

We estimated encounter rates between grazers and prey by modeling the volume of water a swimming grazer will encounter as:

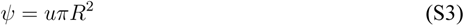

where *u* is the swimming speed and *R* is the perception radius of the grazer. Here, the modeled phytoplankton, *Prochlorococcus* do not swim (Biller *et al*., 2015; Zehr *et al*., 2017). We further account for differences in predator and prey size and encounter efficiency as:

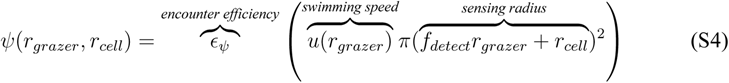

where we assume grazers swimming speed (*µ*m d*^−^*^1^) scales as a function of grazer equivalent spherical radius (*µ*m) using the empirical relation derived by Kiørboe, 2011 as:

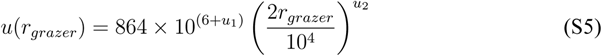

with coefficients *u*_1_ = 0.39 and *u*_2_ = 0.79. We assume grazers detect prey within a detection sensing radius which is proportional to grazer radius with factor *f_detect_* = 3 (Talmy *et al*., 2019b). We also incorporate prey size into the perception radius (Kiørboe, 2008).

##### S2.1.3 Modeling carbon allometry

Ribalet *et al*., 2019 correlate light-scattering to cell size and cell carbon quota using Mie theory. We define cellular elemental quotas (*µ*g per cell) for carbon, *C_S_* as:

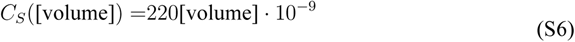

We assume that both infected and susceptible cells have the same elemental quotas. That is *C_I_* = *C_S_*.

For the purposes of converting consumed biomass to grazer abundance we assume grazer quotas in *µ*g per cell for carbon, based on the empirical relationship derived by Menden-Deuer & Lessard, 2000 for dinoflagellates as:

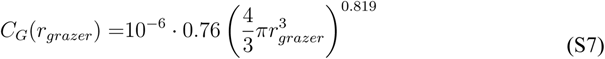

#### S2.2 Viral lysis estimates

##### S2.2.1 iPolony estimates

Estimates for viral-induced lysis based on infected cell data recorded on the cruise using the iPolony method are 0.35-4.8% (Mruwat *et al*., 2021), assuming 1-3 infection cycles occur during the turnover time for *Prochlorococcus* cells.

##### S2.2.2 Viral encounter estimates

Naïve encounter rate estimates are made using the encounter rate theory outlined in equation S1 with baseline parameters assuming that total *Prochlorococcus* loss rates (*ω*) are between 0.3 and 1 per day. Note that this encounter rate estimation assumes instantaneous killing of *Prochlorococcus*. Assuming that the steady state abundance of T4- and T7-like cyanophages during the cruise was *V\** = 3.5 *×* 10^8^ per L, and assuming variation in *E_φ_ ∈* [10*^−^*^5^, 10^0^], *r_virus_ ∈* [20, 50] nm, *r_cell_ ∈* [0.17, 0.49]*µ*m, we find the proportion of *Prochlorococcus* losses via viral-induced lysis can be estimated as:

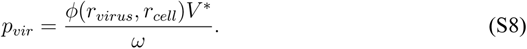

#### S2.3 Grazing estimates

##### S2.3.1 Grazer encounter estimates

Naïve encounter rate estimates are made using the encounter rate theory outlined in equation S4 with baseline parameters assuming that total *Prochlorococcus* loss rates (*ω*) are between 0.3 and 1 per day. Note that this encounter rate estimation assumes instantaneous killing of *Prochlorococcus*. Assuming that the steady state abundance of heterotrophic nanoflagellates during the cruise was *G\** = 2.5 *×* 10^5^ per L, and assuming parameter variation in *E_ψ_ ∈* [10*^−^*^5^, 10^0^], *f_detect_ ∈* [0, 6], *r_grazer_ ∈* [1, 10]*µ*m, *r_cell_ ∈* [0.17, 0.49]*µ*m, we estimate the proportion of *Prochlorococcus* losses via grazing as:

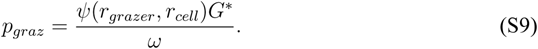

##### S2.3.2 Carbon quota and predator growth rate estimate

Following Connell *et al*., 2020 we use literature values to assume that heterotrophic nanoflagellate growth rate *ξ* is between 0.2 and 1.4 per day (Karayanni *et al*., 2008; Neuer & Cowles, 1994; Verity *et al*., 1993). We can write an approximation for the growth rate of the heterotrophic nanoflagellate population, supposing that they only feed on *Prochlorococcus* cells, as:

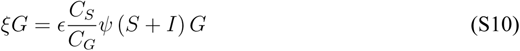

which can be rearranged to estimate the grazing clearance rate as:

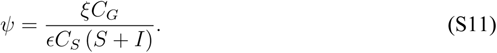

We use a steady state approximation of *P \** = *S\** + *I\** = 1.75 *×* 10^8^ *Prochlorococcus* cells per L and a steady state assumption of *G\** = 2.5 *×* 10^5^ heterotrophic nanoflagellates per L during the cruise. Assuming variation in *E ∈* [0.2, 0.6], *C_S_* and *C_G_* we calculate the proportion of losses attributed to grazing as:

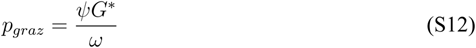

where *ω* represents total *Prochlorococcus* loss rates that we assume are between 0.3 and 1 per day.

##### S2.3.3 FLB estimates

We used Fluorescently Labelled Bacteria (FLB) incubation experiments performed on the cruise (Connell *et al*., 2020) to estimate grazing mortality. These experiments measured the proportion of heterotrophic nanoflagellates containing FLB cells after an hour incubation. The proportion of heterotrophic nanoflagellates containing FLB showed diel variability and varied between 1-25% (Connell *et al*., 2020). In order to use these measurements as the basis for estimating grazing mortality we propose that the population dynamics of FLB cells within incubation experiments can be modeled as:

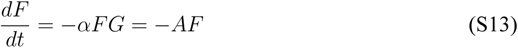

where F is the abundance of FLB cells, *α* is the clearance rate of heterotrophic nanoflagellates feeding on FLB and *A* represents the grazing rate by heterotrophic nanoflagellates. Assuming that heterotrophic nanoflagellates graze on *Prochlorococcus* and FLB at the same rate, and assuming a constant heterotrophic nanoflagellate population, *G*, of 2*×*10^5^ cells/L allows a grazing rate *A* to be computed as:

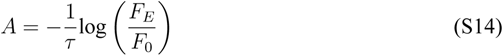

where *τ* is the experiment length (1 hour), the initial FLB abundance is *F*_0_ = 10^8^ cells per L, and the final FLB abundance is *F_E_*. Assuming the observed proportion (0.01-0.25) of heterotrophic nanoflagellates containing FLB (*Y*) consume *X* FLB on average during the incubation period, we can calculate the final FLB abundance as:

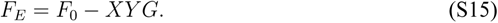

We assume that *X* varies between 1 and 25, but, it is unclear exactly how many FLB were eaten per grazer during the 1 hour incubation. We assume that estimated grazing rates on FLB are equivalent to grazing rates on *Prochlorococcus*. To convert to mortality, we assume that total *Prochlorococcus* loss rates, *ω* are between 0.3 and 1 per day and calculate proportional losses by grazing as:

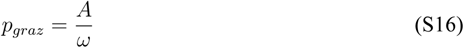

### S3 ECLIP model and parameter inference

For completeness, and to help the reader, we repeat our descriptions of the ECLIP model in the supplementary text.

#### S3.1 ECLIP: Ecological Community driven by Light, with Infection of Phytoplankton

The ECLIP model represents *Prochlorococcus* division and death where division is light-driven (where cell division is expected to occur at night (Binder & DuRand, 2002; Ribalet *et al*., 2015)) and death is controlled by viral lysis, grazing, and other density-dependent factors (Figure 1a). The *Prochlorococcus* population is structured by two states of infection: cells that are susceptible to viral infection (*S*) and cells that are infected (*I*) by viruses (*V*). Grazers (*G*) feed indiscriminately on both *S* and *I* classes. The dynamics of *S*, *I*, *V* and *G* numerical abundances over time are described by the following system:

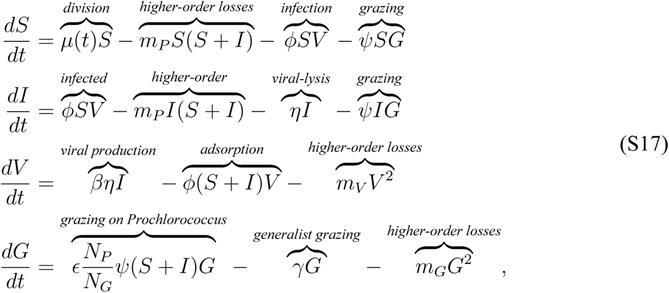

Where

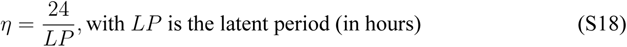

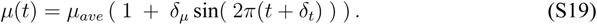

*Prochlorococcus* have a diel-driven population division rate *µ*(*t*) whose proportional amplitude and phase are set by parameters *δ_µ_* and *δ_t_*, and *t* = 0 represents 06:00:00 local time (see Figure S1). *Prochlorococcus* have a nonlinear loss rate, *m_P_*, dependent on total phytoplankton population size to implicitly represent niche competition (Berube *et al*., 2019). Viruses infect susceptible *Prochlorococcus* at a rate *φ* and create *β* new virions that are released into the environment upon cellular lysis, which occurs after an infection latent period of 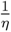. Grazing upon *Prochlorococcus* is non-preferential with respect to infection status and occurs at a rate *ψ* with a Gross Growth Efficiency (GGE) *E* proportional to the fraction of nitrogen contents in a *Prochlorococcus* cell (*N_P_* = 5.01 *×* 10*^−^*^9^*µ*g N cell*^−^*^1^) and a grazer (*N_G_* = 6.53 *×* 10*^−^*^6^*µ*g N cell*^−^*^1^). We introduce *γ* as a parameter to denote whether grazers act as specialists (*γ* = 0) or generalists (*γ >* 0), where the term represents net additional gains to the grazer from non-*Prochlorococcus* prey sources after accounting for respiratory costs. Generalist strategies may include ingesting other phytoplankton, heterotrophic bacteria, or other grazers through intraguild predation. Grazer and viral losses are both characterized by a nonlinear loss term to avoid structurally biasing the model to favour one of these types of *Prochlorococcus* predators (Talmy *et al*., 2019a) and to avoid competitive exclusion. A full list of parameters are shown in Table S2.

##### S3.1.1 Defining the specialism-generalism gradient

We define the degree of generalism (*d*_generalism_) as the ratio of growth performed by grazers derived from other sources relative to that derived from both other sources and from consuming *Prochlorococcus*:

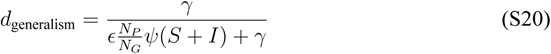

**Figure S1:**
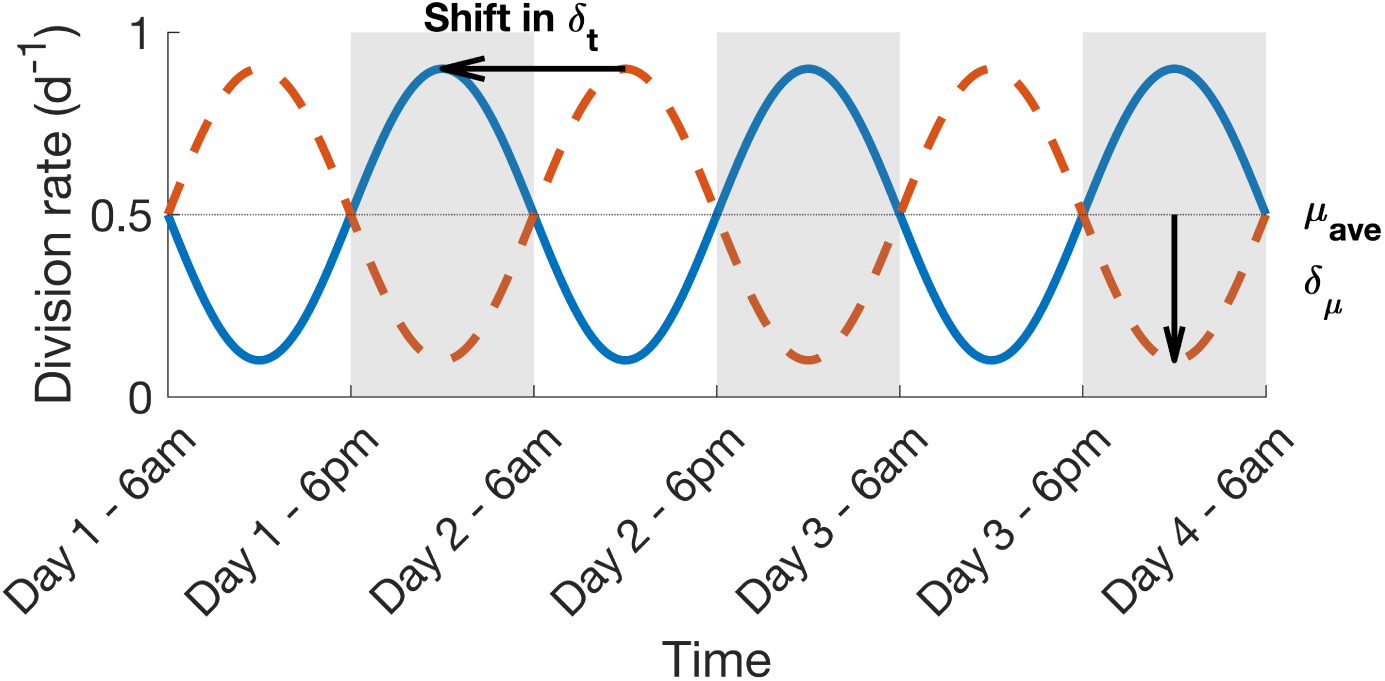
Diel-driven division rate. Solid blue and dashed orange lines are example division rate curves to graphically explain how the different parameters used in the diel-driven division rate function in ECLIP alter the model division rate. The parameters for division rate are represented by the average division rate over 1 day, *µ_ave_*, the amplitude of the sinusoidal oscillations, *δ_µ_*, and the phase of the sinusoidal signal, *δ_t_*. The two curves differ in their value of *δ_t_*; the orange curve has *δ_t_*= 0, while *δ_t_* = 0.5 days for the blue curve.

##### S3.1.2 Model inferred mortality

Total *Prochlorococcus* loss rates are defined as:

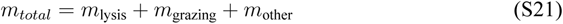

where:

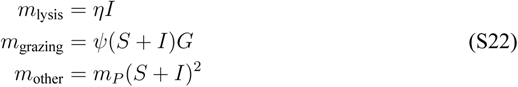

We defined the proportion of each mortality processes as follows:

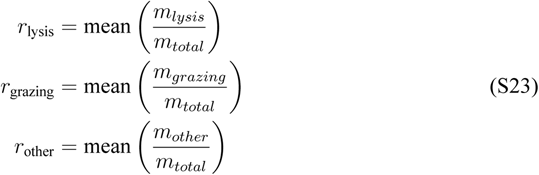

##### S3.1.3 Diel adsorption rates

We incorporated a diel adsorption rate into our analyses in Figure 6 to see if this would better represent the observed population dynamics of infected *Prochlorococcus* cells. We did this by modulating the average inferred adsorption rate with a time dependent step function such that adsorption rates are highest at dusk, and lowest at dawn - consistent with lab experiments and associated modeling between cyanophage and *Prochlorococcus* (Demory *et al*., 2020; Liu *et al*., 2019).

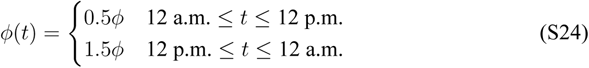

**Table S2:**
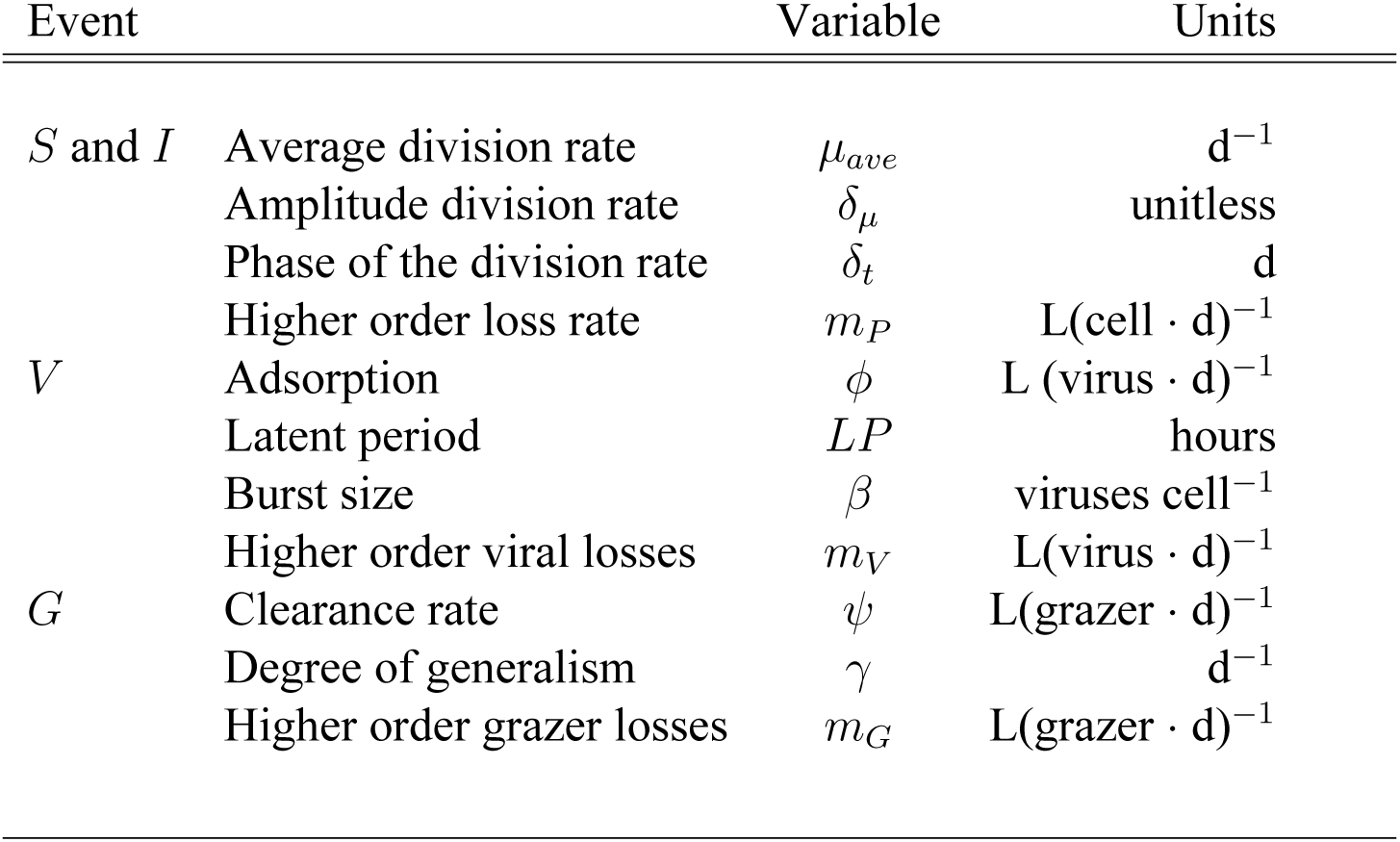
Parameters of the the ECLIP model

#### S3.2 Model-data integration

##### S3.2.1 General parameter inference implementation

We determined the parameter sets for the ECLIP model that optimized the fit of the model dynamics to field measurements by using Markov Chain Monte Carlo (MCMC) implemented in the probabilistic inference package Turing (Ge *et al*., 2018) in the Julia language (Bezanson *et al*., 2017). MCMC is a class of Bayesian inference algorithms that aims to infer the probability distribution of the model parameters given the model equations, environmental data and prior beliefs (Lambert, 2018). We used the No-U-Turn Sampler (NUTS) implemented in Turing to sample the posterior distributions (Hoffman, Gelman, *et al*., 2014). Classic Hamiltonian Monte Carlo (HMC) algorithms are sensitive to the number of steps of the simulation reducing the performance of the algorithm in high-dimension problems. NUTS is a performant alternative algorithm for MCMC that eliminates the need to set the number of steps, allowing better performance for high-dimensional parameter space and non-linear dynamical models (Hoffman, Gelman, *et al*., 2014).

We implemented the parameter estimation in two steps. First, we estimated the three parameter posterior distributions (*µ_ave_*, *δ_µ_* and *δ_t_*) for the division rate function (see equation S19) using a model-inferred *Prochlorococcus* division rate estimated for the same oceanographic campaign by (Mattern *et al*., 2022). In a second step, we used the division parameter posteriors as priors for another parameter inference to estimate the whole set of model parameters (*µ_ave_*, *δ_µ_*, *δ_t_*, *m_P_*, *m_V_*, *m_G_*, *φ*, *β*, *LP*, and *ψ*) and the abundance measurements for total and infected *Prochlorococcus* cells, cyanophages and heterotrophic grazers. We repeated this second step six times by fixing the generalism term at *γ* = 0, 0.01, 0.05, 0.1, 0.2 and 0.5 respectively, to account to different levels of non-*Prochlorococcus* contributions to grazing.

For each implementation, we ran 4 MCMC chains for 4000 iterations with a 2000 iteration warm-up period (total iterations = 6000) and a target acceptance ratio of 0.65, allowing the algorithm to reach a stationary state. For the model *γ* = 0.5, we obtained chains with high autocorrelation. To decrease the autocorrelation we ran 16 independent chains of 4000 iterations and combined them in four longer chains of 16000 iterations. We then thinned these longer chains (Link & Eaton, 2012) by sampling every 4 iterations to obtain 4 MCMC chains of 4000 iterations with lower autocorrelation.

##### S3.2.2 Chain convergence analysis

To assess the convergence of the MCMC chains, we computed different indicators for each implementation (Gelman *et al*., 1995). For each parameter, the inter-variability of the chains was computed as the 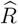. When the set of chains converged to the same posterior distribution, 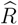 should be centered around 1 (with upper bound of 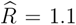), suggesting good convergence (Figure S5). We computed the ratio of effective sample size (*N_eff_*) to the total iteration size (*N* = 4000) to search for potential sampling issues. Low ratio 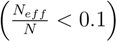, indicates potential sampling problems with higher chain auto-correlation, suggesting divergence (Figure S5). Additionally, we also computed auto-correlation for each parameter chains of the ECLIP model (Figure S8).

##### S3.2.3 Division function parameter inference

We fitted our division function (eq. S19) to the timeseries of division rates estimated by integrating *in situ* flow cytometry measurements with the pmb size-structured *Prochlorococcus* model described in Mattern *et al*., 2022. Daily division rates were estimated using a size-structured matrix population model that mechanistically describes changes in microbial cell size distribution over the day-night cycle (Mattern *et al*., 2022). In brief, the model assumes that changes of the cell size distribution is driven by three interconnected biological processes: carbon fixation via photosynthesis, carbon loss via respiration and exudation and cell division. We select the pmb model with a power-law size dependence on carbon fixation since there is strong evidence of an allometric growth rate in natural populations of *Prochlorococcus* (Casey *et al*., 2019). The model was fit to a logarithmically spaced discrete cell size distribution of *Prochlorococcus* provided by SeaFlow, using the same priors of model parameters used in Mattern *et al*., 2022.

This implementation helped us to refine the division priors for the whole model parameter estimation step. We used wide uniform priors for the three parameters of the division function: *µ_ave_*, *δ_µ_* and *δ_t_* (Figure S2, Table S3). The 4 chains converged (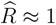, with no auto-correlation), to narrower distributions centered around *µ_ave_* = 0.48 *d^−^*^1^, *δ_µ_* = 0.75 *d^−^*^1^ and *δ_t_* = 0.83 *d*. Our division function predicted well the division rates from Mattern *et al*., 2022 (Figure S3).

**Figure S2:**
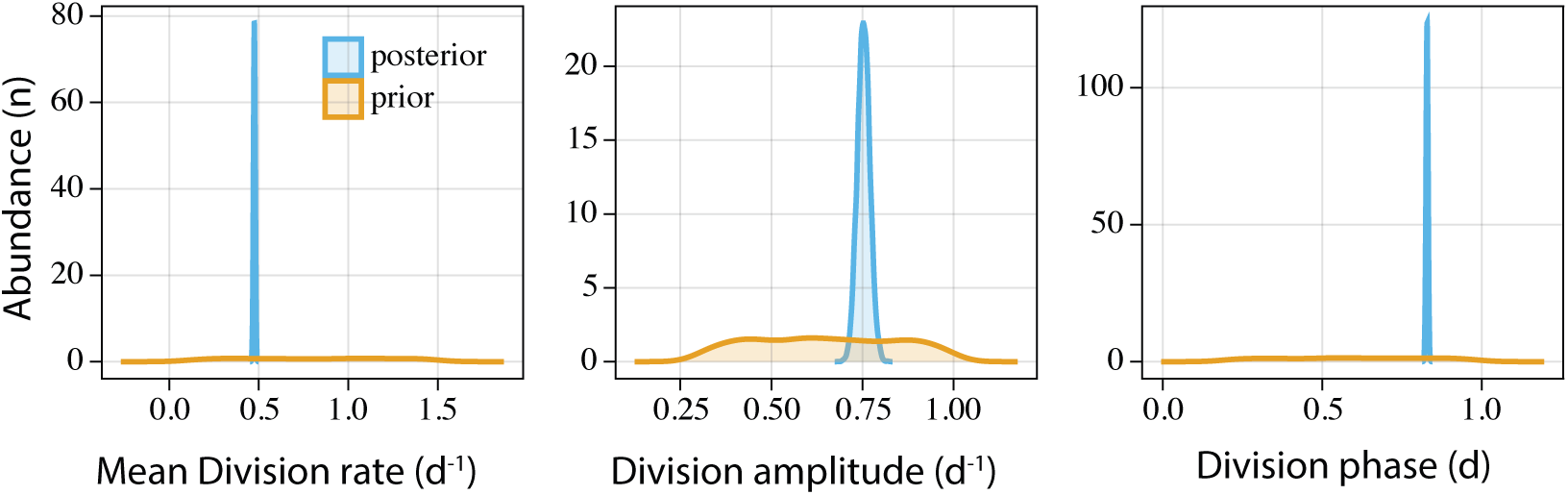
Priors and posteriors distribution for the division function parameters. We used wide uniform priors (orange) allowing to search a wide space for each parameter. Posteriors (blue) had narrower distributions that helped us to reduce the parameter range for the whole parameter inference step.

**Figure S3:**
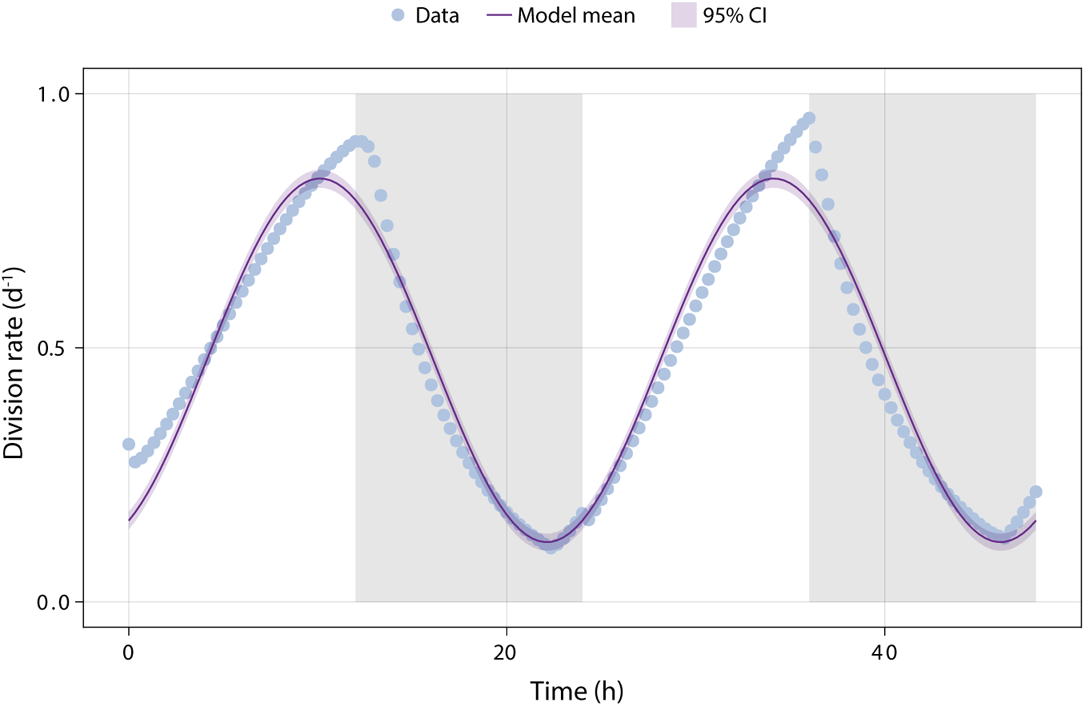
Division function fits. Our periodic function (purple) well represented the estimated division rate (blue dot) from (Mattern *et al*., 2022). Gray vertical shaded areas represent the night periods.

##### S3.2.4 ECLIP parameters inference implementation

To estimate the whole parameter set for the ECLIP model, we used environmental abundance data from the SCOPE HOE-Legacy 2A cruise (see Empirical data section, 2.3 in the main text, for more information). Specifically, we used total and infected *Prochlorococcus* cells, free cyanophage and heterotrophic bacteria. We inferred parameters of the ECLIP model for six levels of generalism: *γ* = 0, 0.01, 0.05, 0.1, 0.2, and 0.5 day*^−^*^1^. For each implementation, we used the same priors for model parameters and log-likelihood (see next section for more information on the priors). Comparison between posteriors of the 6 models and the priors is shown in Figure S4.

**Figure S4:**
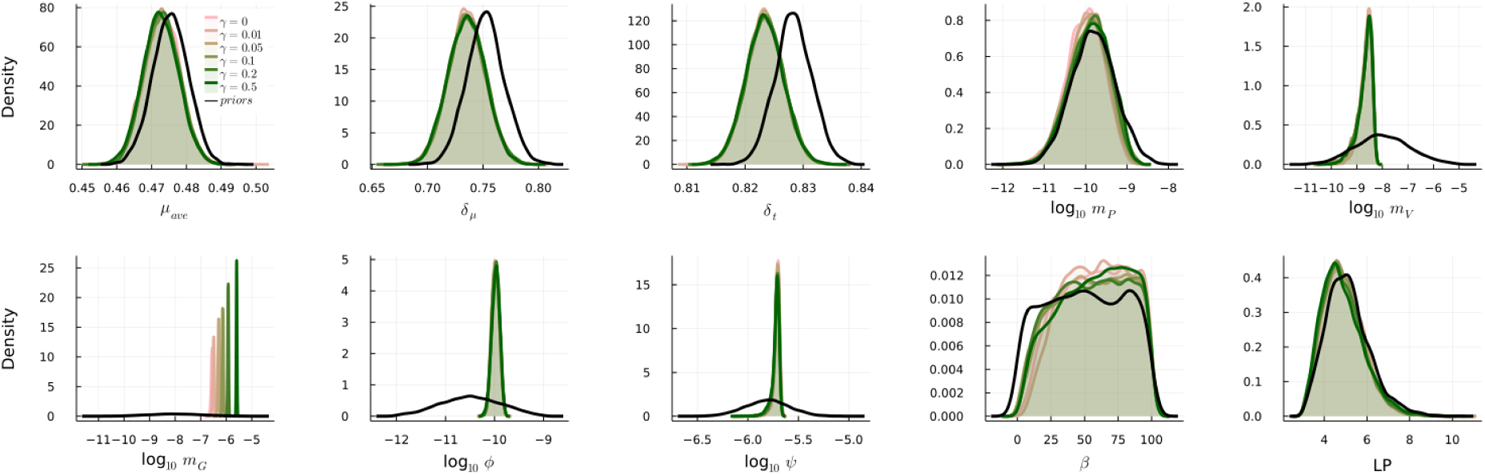
Comparison of parameter prior and posterior estimates in ECLIP. Prior distributions are represented in solid black lines, while posteriors are represented in colored solid lines (from pink *γ* = 0 d*^−^*^1^ to dark green *γ* = 0.5 d*^−^*^1^).

**Table S3:**
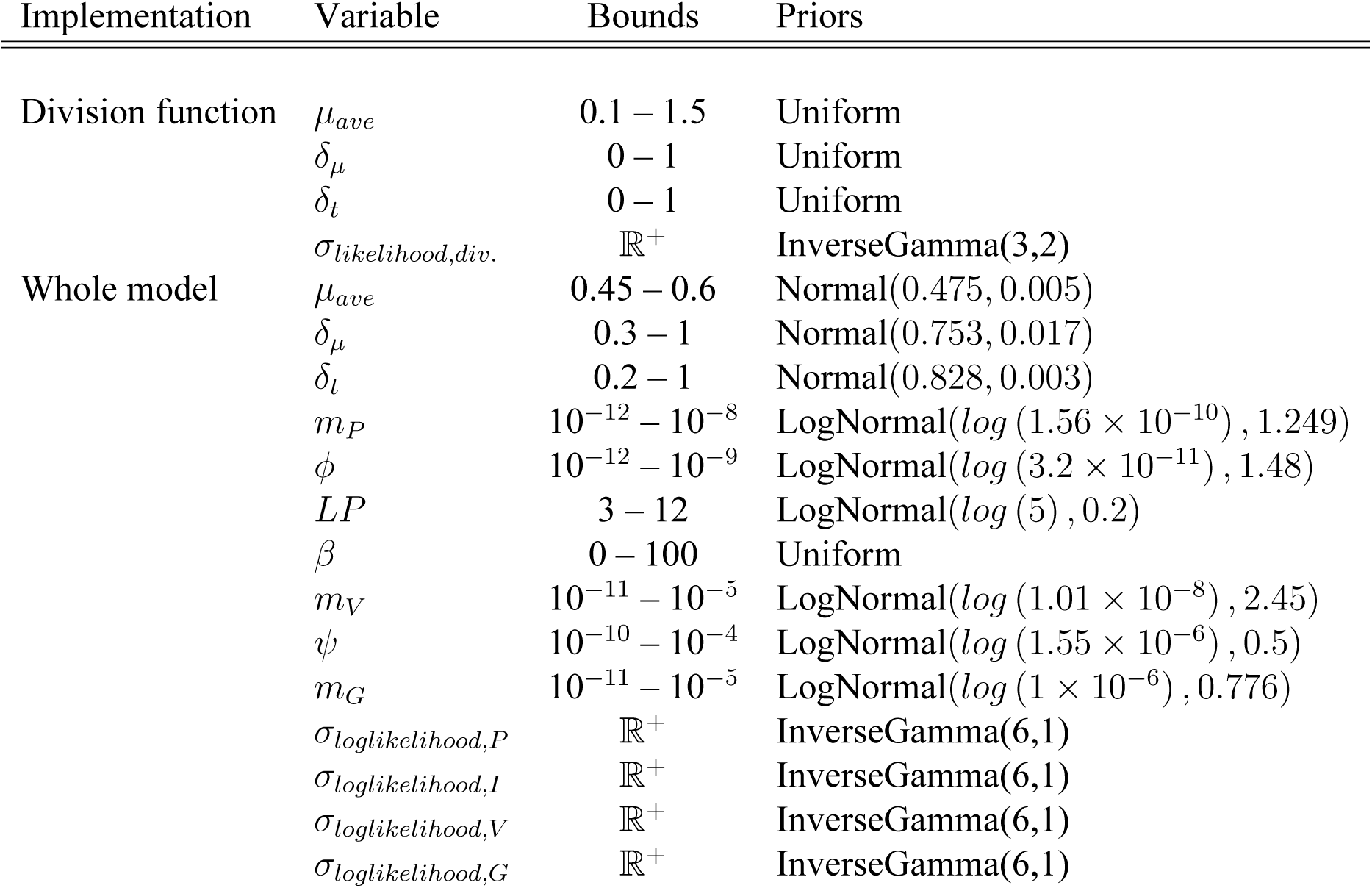
Priors and parameter ranges explored for the parameter inferences

##### S3.2.5 Priors and parameter ranges

We defined priors for both inference implementations, for each of the model parameters and the standard deviation *σ* of the likelihoods (Table S3). The ECLIP model is prone to identifiability problems due to parameter correlations, possibly leading to performance and convergence issue of the MCMC algorithm. Therefore, we weighted higher probability space for some parameters by setting Normal or log-Normal priors instead of Uniform priors for the majority of the parameters, while setting wide variance for each prior, to explore the larger parameter space as possible given ecological realistic ranges. Additionally, for the whole implementation, we fixed three parameters to reduce parameter identifiability issues: the gross growth efficiency (*E* = 0.3, (Straile, 1997)), and the fraction of Nitrogen content for *Prochlorococcus* (*N_P_* = 5.01*×*10*^−^*^9^*µ*g N cell*^−^*^1^) and grazers (*N_G_* = 6.53 *×* 10*^−^*^6^*µ*g N cell*^−^*^1^).

### S3.3 Adding diel-dependent adsorption rates

In Figure S11 we added step-wise diel-dependent adsorption rates to the core ECLIP models, as described in S3.1.3.

**Figure S5:**
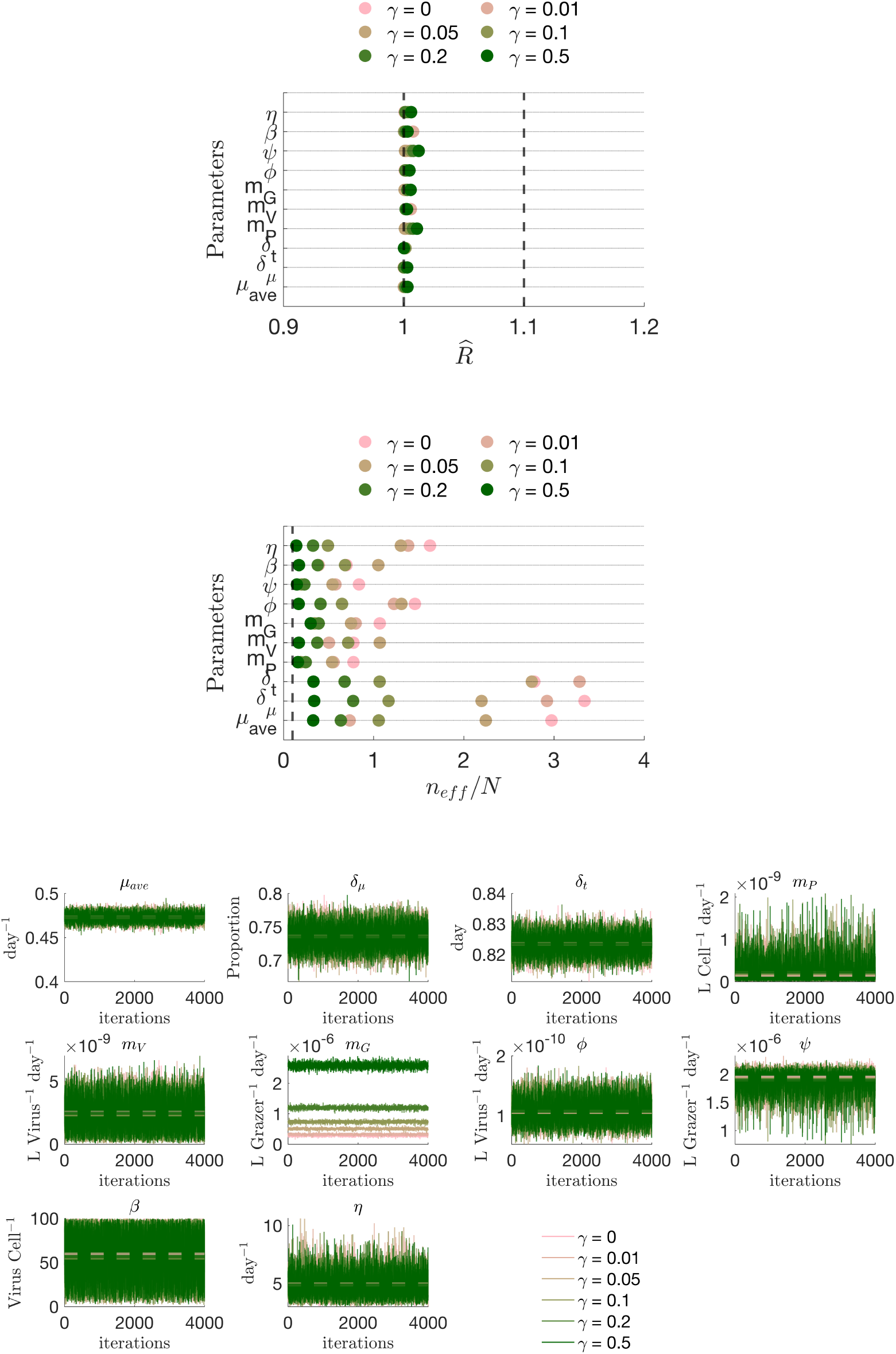
Assessing MCMC convergence and efficiency for ECLIP models. (a) *R* convergence diagnostics. (b) Ratio of effective sample size to sample size. (c) Visualizing MCMC parameter chains for each of the ECLIP models (note density plots from chains are shown in Figure 3c). Full details of parameter bounds are shown in Table S2.

**Figure S6:**
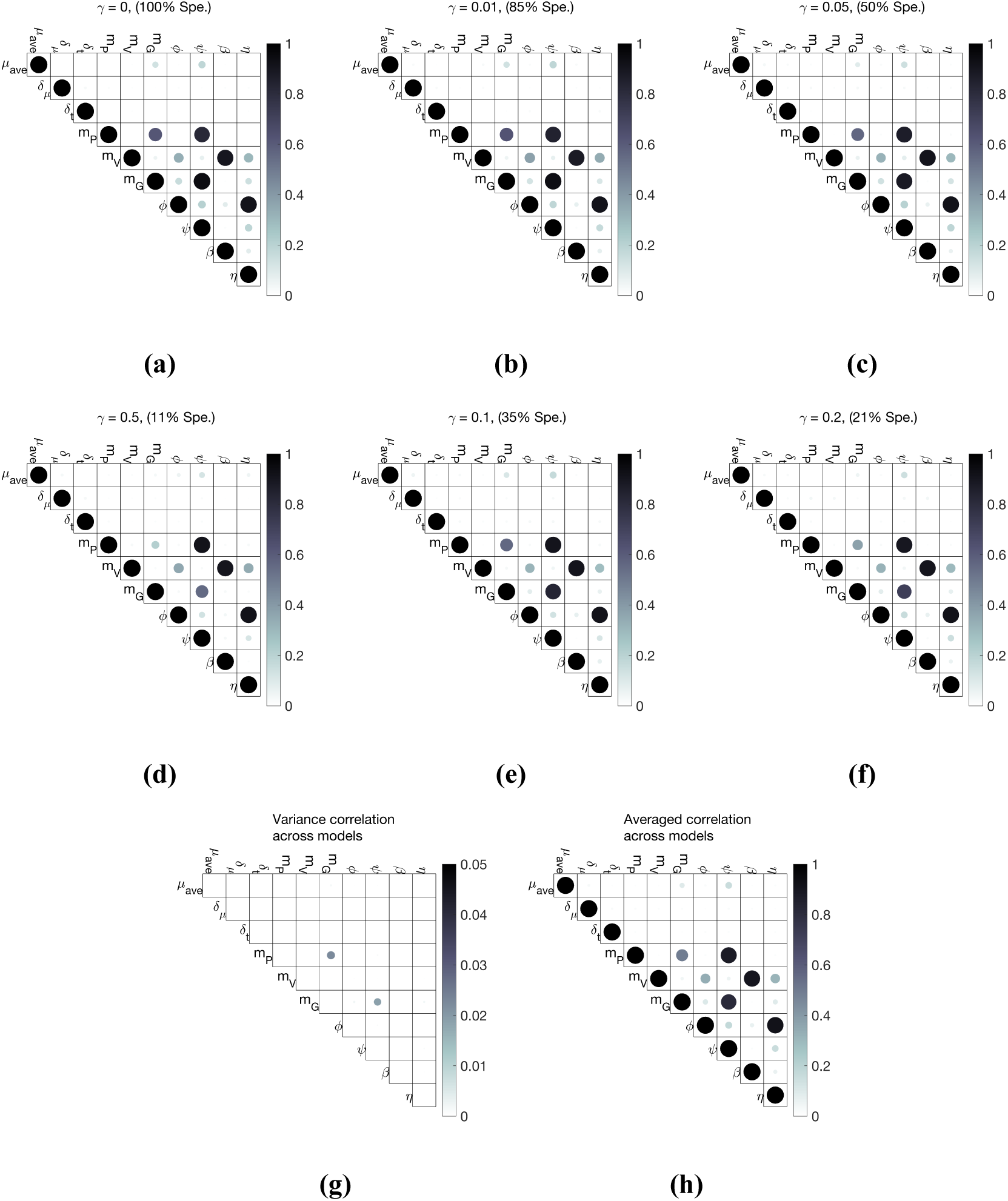
Identified parameter correlation in ECLIP models. Pearson correlation between parameters in MCMC chains for models with: (a) *γ* = 0 d*^−^*^1^, (b) *γ* = 0.01 d*^−^*^1^, (c) *γ* = 0.05 d*^−^*^1^, (d) *γ* = 0.1 d*^−^*^1^, (e) *γ* = 0.2 d*^−^*^1^, (f) *γ* = 0.5 d*^−^*^1^. (g) Pearson correlation between parameters across all MCMC chains together. (h) Average of the assessed correlations between parameters across grazer strategy models (a-f).

**Figure S7:**
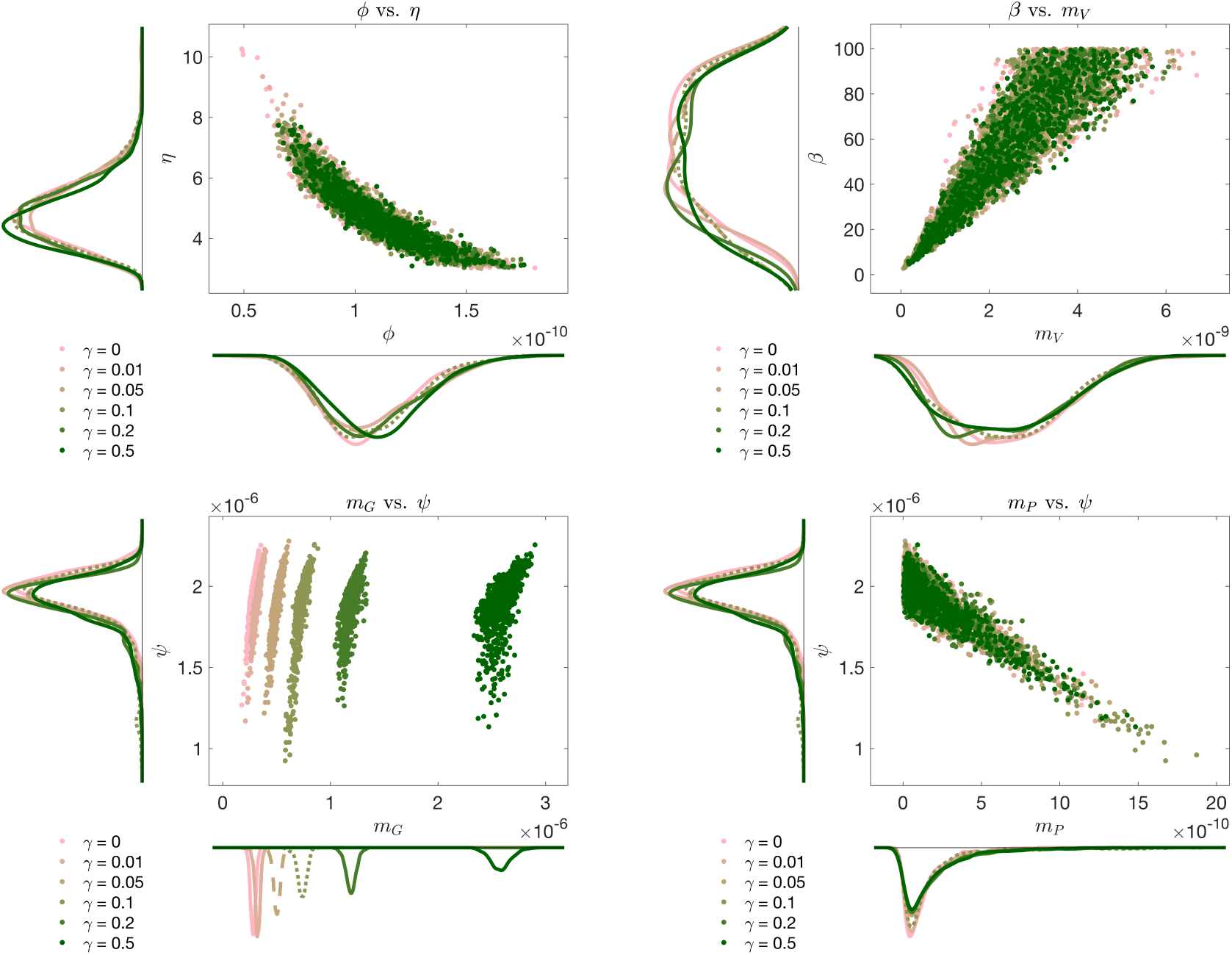
Examples of parameter covariance in ECLIP. Scatterplots of highly correlated ecological parameters: (a) adsorption rate (*φ*) vs lysis rate (*η*), (b) burst size (*β*) vs. viral higher order loss (*m_V_*), (c) grazer higher order loss (*m_G_*) vs. grazing rate (*ψ*), (d) *Prochlorococcus* higher order loss (*m_P_*) vs. grazing rate (*ψ*).

**Figure S8:**
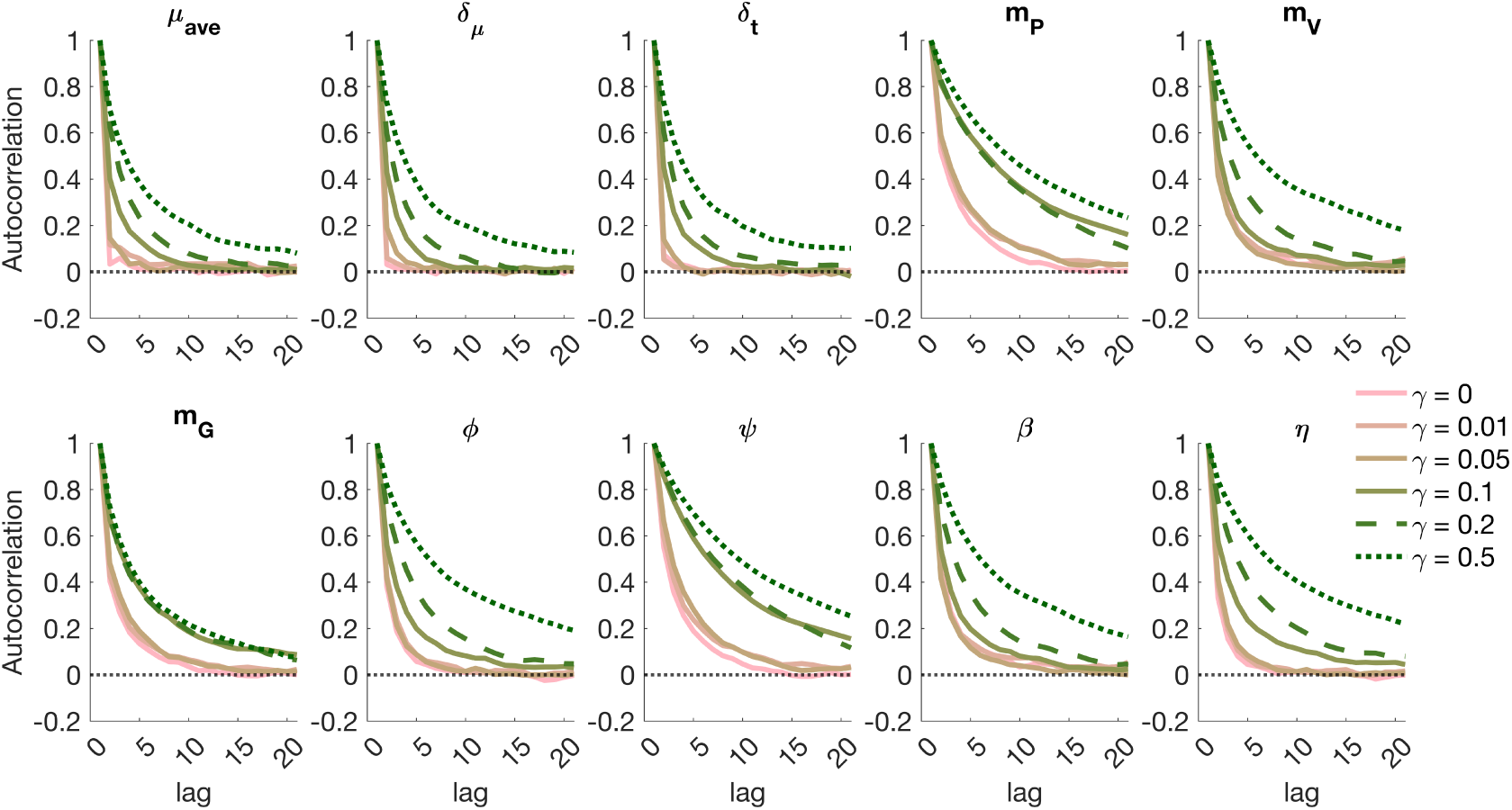
Parameter autocorrelation across ECLIP models. Autocorrelation was calculated as the cross-correlation of the chain with itself. Strong autocorrelation is leading to patterns or periodicity in the chain and suggest bad convergence. Our 6 models have relatively low autocorrelation for each parameter suggesting convergence.

**Figure S9:**
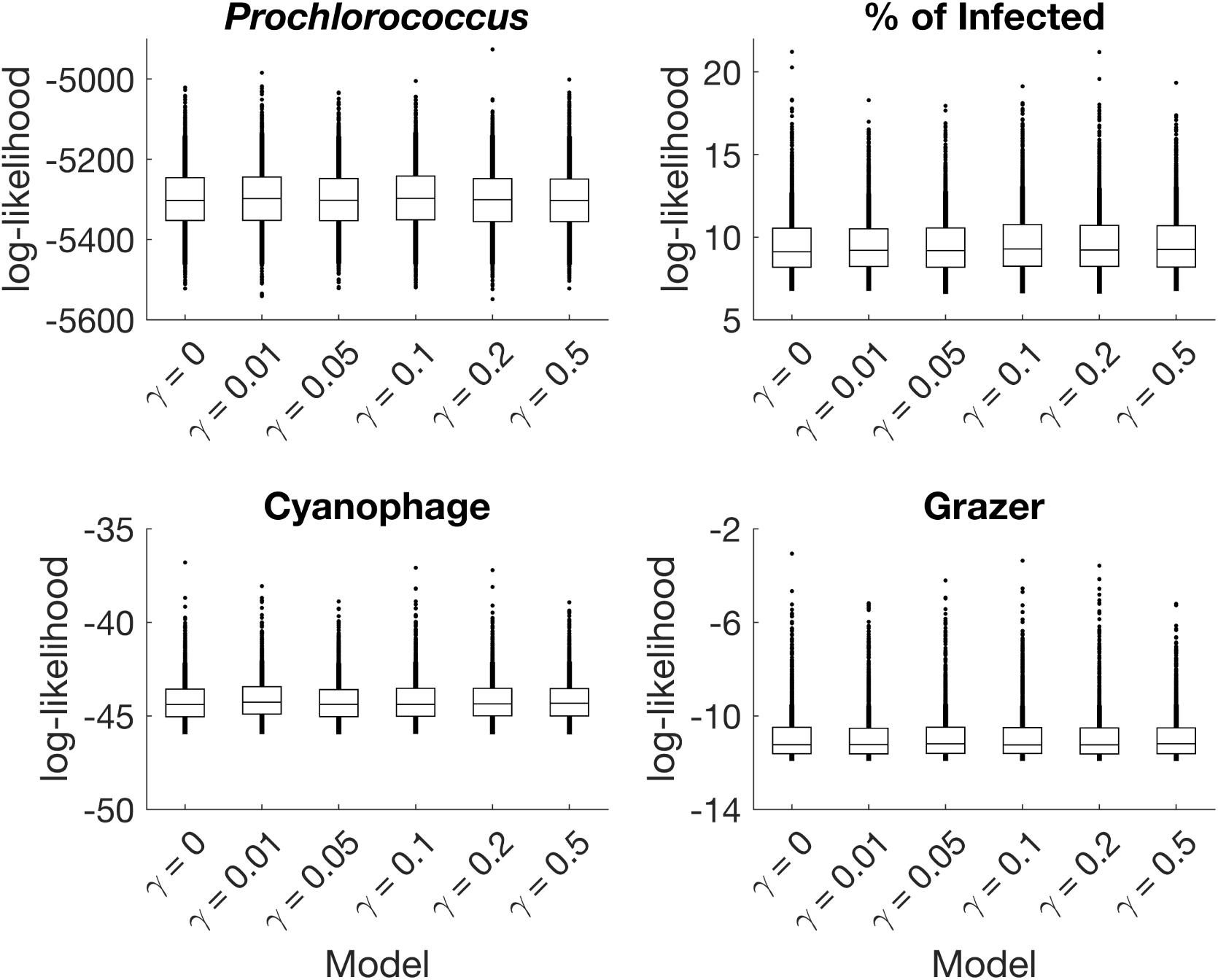
Negative log-likelihood for *Prochlorococcus*, % of infected *Prochlorococcus*, cyanophage and grazer across the 6 models. The negative log-likelihood function was calculated, for each chain step, as 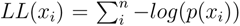, where *p*(*x_i_*) is the model probability density function following a log-normal distribution and *n* the number of observations. The best model has the minimum *LL*. *LL* can only be compared between models for the same observations channels (*Prochlorococcus*, % of infected, Cyanophage or Grazer) and not between channels. The 6 models have the same ability to reproduce the observations for each chanels. Boxplots show the variability of the *LL* across the MCMC chain.

**Figure S10:**
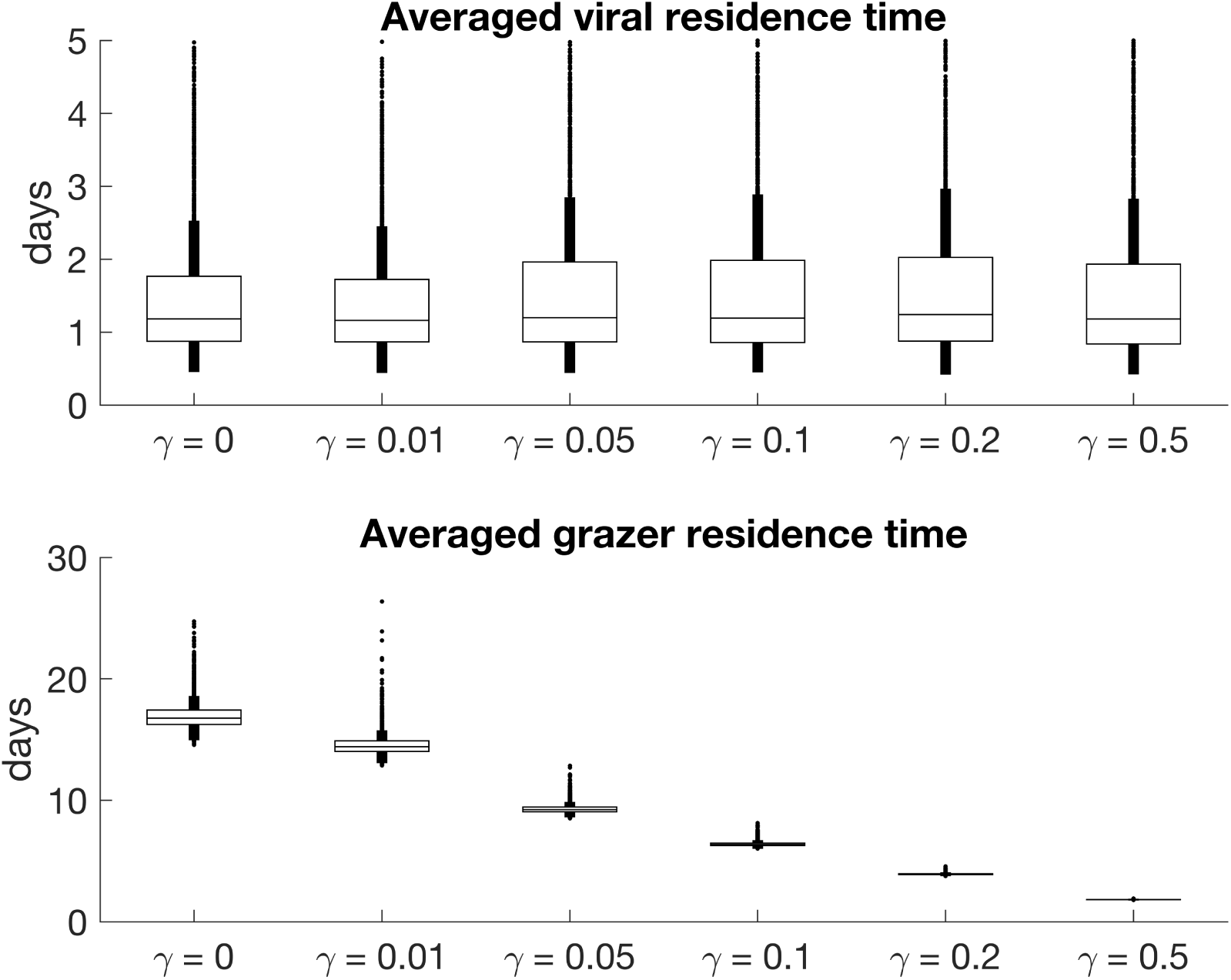
Averaged residence time of cyanophage (upper panel) and grazer (lower panel). Viral and grazer residence times were calculated, for each chain step, as *T_V_* = *mean*(1/(*m_V_ · V* (*t*)) and *T_G_* = *mean*(1/(*m_G_ · G*(*t*)) respectively. Median virus residence times are similar across the 6 models and range from 1.16 to 1.22 days, whereas median grazer residence times decreased from model 16.8 days (model γ = 0) to 1.8 days (model γ = 0.5). Boxplots show the variability of the log-likelihood across the MCMC chain.

**Figure S11:**
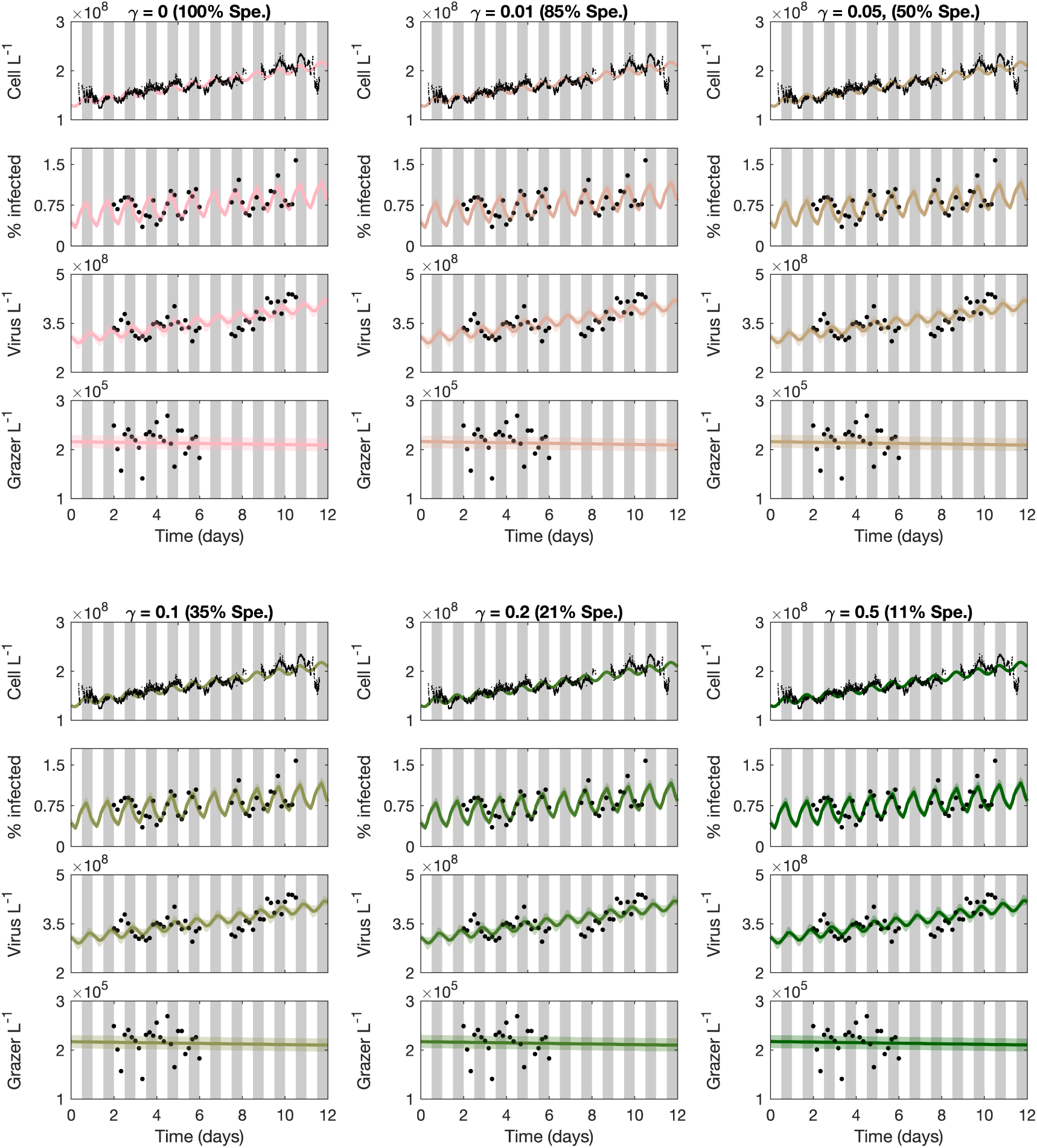
Models with diel adsorption rates across the specialism-generalism gradient fit empirical data. ECLIP models are compared against empirical data in black. Model lines represent the median MCMC solution within 95% CI range found by the converged chains, shown as bands with colours representing the choice of *γ*. Data signals include *Prochlorococcus* cell abundances (top), the percentage of infected *Prochlorococcus* cells, the abundance of free viruses and the abundance of heterotrophic nanoflagellate grazers (bottom). The models were fitted against detrended data; for visualization we have added these trends to the model solutions. Grey bars indicate nighttime. Model solutions with: (a) *γ* = 0 (grazers act as specialists), (b) *γ* = 0.01, (c) *γ* = 0.05, (d) *γ* = 0.1, (e) *γ* = 0.2, (f) *γ* = 0.5 day*^−^*^1^. The degree of grazer specialism (Spe.) is shown in parentheses above each subplot.

### S4 Potential mechanisms to explain other losses of *Prochlorococcus*

### Undercharacterized ecological interactions

Mixotrophic nanoflagellates contribute to *Prochlorococcus* grazing, with measurements suggesting smaller contribution relative to the heterotrophic nanoflagellates (Connell *et al*., 2020; Li *et al*., 2021) – but, still likely significant (Li *et al*., 2022; Zubkov & Tarran, 2008). Evaluating *in situ* mixotrophy is an ongoing challenge. Unknown viruses not quantified by the iPolony method (e.g., cyanophage without an identifiable DNA polymerase gene - see Cai *et al*., 2022), and other grazer types including larger consumers may contribute to ‘other mortality’. Additionally, in ECLIP viral-induced and grazing-induced losses utilise a contact-driven model, analogous to ‘Type I’ functional responses. Mechanistic changes in functional responses and/or responses to light may drive distinct interaction rates (and aggregate mortality) even given the same set of viruses and grazers.

### Aggregation and sinking of picoplankton

Picoplankton (and their viruses) are implicated as important contributors to export in large scale analyses e.g., (Guidi *et al*., 2016; Richardson & Jackson, 2007), though Guidi *et al*., 2016 suggest *Synechococcus* rather than *Prochlorococcus* abundances appear correlated to export. Less is known about microscale processes leading to picoplankton export contributions – *Prochlorococcus* cells can be sustained in laboratory culture for month-long experiments, suggesting limited cell sedimentation. Generally, conceptual models of ocean ecology don’t include sinking out of picoplanktonic populations – assuming it is inconsequential (Richardson, 2019). However, particle aggregation could be stimulated via sloppy feeding or viral lysate (Sullivan *et al*., 2017; Talmy *et al*., 2019a). As viral lysate from picocyanobacteria is labile (Xiao *et al*., 2021; Zhao *et al*., 2019; Zheng *et al*., 2021) aggregation and sinking could be stimulated indirectly via heterotrophic bacterial growth. Particle attachment and aggregation could also be stimulated by TEP (Transparent exopolymer particles), produced in xenic *Prochlorococcus* cultures (Cruz & Neuer, 2019; Iuculano *et al*., 2017). TEP production can be stimulated by high light intensity, characteristic of the surface ocean, and its production appears linked to loss rates (Iuculano *et al*., 2017). Picoplankton export could also be attributed to mineral ballasting (Richardson, 2019) – both *Synechococcus* (Baines *et al*., 2012; Krause *et al*., 2017) and *Prochlorococcus* (Krause *et al*., 2017; Wei *et al*., 2021) have been suggested to accumulate biogenic silica.

### Physiological stress(es)

Other cyanobacterial loss mechanisms include physiological stress, induced for example, by high-level irradiance associated with ultraviolet radiation (Llabrés *et al*., 2011; Mella-Flores *et al*., 2012) and refraction of light through surface waves that could lead to photodamage via the flashing effect (Demory *et al*., 2018; Stramska & Dickey, 1998). Other abiotic factors including nutrient deficiency (Kulk *et al*., 2013), metal toxicity (Mann *et al*., 2002; Mann & Chisholm, 2000; Sarker *et al*., 2021), and thermal variations (Baker & Geider, 2021; Pittera *et al*., 2014) can also contribute to increased stress, though we do not expect significant thermal variations in the NPSG (mean seasonal changes are +/- 3°C (23.5-26.5 °C)). Stressors often results in the generation of reactive oxygen species, that cause oxidative stress and can cause DNA damage. Reactive oxygen species are also used as a signalling pathway for programmed cell death in photosynthetic microbes (Bidle, 2015, 2016; Vardi *et al*., 2006), though evidence is lacking for a programmed cell death pathway in *Prochlorococcus* and sympatric heterotrophic bacteria are thought to alleviate this stress (Morris *et al*., 2011, 2008). We note evidence is also lacking for toxin-antitoxin systems, common across bacteria and archaea which could lead to cell loss, in *Prochlorococcus* (Fucich & Chen, 2020).

### Population heterogeneity

Microbes experience senescence and aging, leading to intracellular accumulation of damage through their life cycle (Moger-Reischer & Lennon, 2019), which may lead to asymmetric division (Franklin, 2013). Unlike other microorganisms, *Prochlorococcus* has no resting stages and relies on heterotrophic bacteria to survive nutrient starvation in cultures (Roth-Rosenberg *et al*., 2020), though the heterotrophic bacteria they form associations with *in situ* differ from those found in culture which tend to be copiotrophic (Kearney *et al*., 2021). *Prochlorococcus* populations are combinations of ecotypes which respond differently to environmental stressors (Johnson *et al*., 2006; Martiny *et al*., 2006; Moore & Chisholm, 1999; Thompson *et al*., 2011; Tolonen *et al*., 2006; Zinser *et al*., 2007). Heterogeneity within the *Prochlorococcus* population, potentially via vertical displacement, may mean there are differing mortality responses at the individual or strain-level vs. at the scale of total population. Additionally, competition between ecotypes and with other phytoplankton in the same niche (like *Synechococcus* or picoeukaryotes) may also increase stresses, especially in a nutrient limited oligotrophic environment like the NPSG.

## References

Acevedo-Trejos, E., Brandt, G., Smith, S. L. & Merico, A. (2016). PhytoSFDM version 1.0. 0: phytoplankton size and functional diversity model. Geoscientific Model Development, 9, 4071–4085.

Aguilera, A., Klemencic, M., Sueldo, D. J., Rzymski, P., Giannuzzi, L. & Martin, M. V. (2021). Cell death in Cyanobacteria: current understanding and recommendations for a consensus on its nomenclature. Frontiers in Microbiology, 12, 631654.

Arias, A., Saiz, E. & Calbet, A. (2020). Towards an Understanding of Diel Feeding Rhythms in Marine Protists: Consequences of Light Manipulation. Microbial Ecology, 79, 64–72.

Ashkezari, M. D., Hagen, N. R., Denholtz, M., Neang, A., Burns, T. C., Morales, R. L., et al. (2021). Simons collaborative marine atlas project (Simons CMAP): an open-source portal to share, visualize, and analyze ocean data. Limnology and Oceanography: Methods, 19, 488–496.

Aylward, F. O., Boeuf, D., Mende, D. R., Wood-Charlson, E. M., Vislova, A., Eppley, J. M., et al. (2017). Diel cycling and long-term persistence of viruses in the ocean’s euphotic zone. Proceedings of the National Academy of Sciences, 114, 11446–11451.

Azam, F., Fenchel, T., Field, J. G., Gray, J. S., Meyer-Reil, L.-A. & Thingstad, F. (1983). The ecological role of water-column microbes in the sea. Marine ecology progress series, 257–263.

Baran, N., Goldin, S., Maidanik, I. & Lindell, D. (2018). Quantification of diverse virus populations in the environment using the polony method. Nature Microbiology, 3, 62–72.

Becker, K. W., Collins, J. R., Durham, B. P., Groussman, R. D., White, A. E., Fredricks, H. F., et al. (2018). Daily changes in phytoplankton lipidomes reveal mechanisms of energy storage in the open ocean. Nature Communications, 9, 5179.

Beckett, S. J., Demory, D., Coenen, A. R., Casey, J. R., Dugenne, M., Follett, C. L., et al. (2021). Diel population dynamics and mortality of Prochlorococcus in the North Pacific Subtropical Gyre. bioRxiv,

Beckett, S. J., Demory, D., Coenen, A. R., Casey, J. R., Dugenne, M., Follett, C. L., et al. (2023). Code for: Disentangling top-down drivers of mortality underlying diel population dynamics of Prochlorococcus in the North Pacific Subtropical Gyre. Version ECLIP2.

Beckett, S. J. & Weitz, J. S. (2017). Disentangling niche competition from grazing mortality in phytoplankton dilution experiments. PLOS ONE, 12, e0177517.

Beckett, S. J. & Weitz, J. S. (2018). The effect of strain level diversity on robust inference of virus-induced mortality of phytoplankton. Frontiers in Microbiology, 9, 1850.

Berube, P. M., Rasmussen, A., Braakman, R., Stepanauskas, R. & Chisholm, S. W. (2019). Emergence of trait variability through the lens of nitrogen assimilation in *Prochlorococcus*. elife, 8, e41043.

Bezanson, J., Edelman, A., Karpinski, S. & Shah, V. B. (2017). Julia: A fresh approach to numerical computing. SIAM review, 59, 65–98.

Binder, B. J. & DuRand, M. D. (2002). Diel cycles in surface waters of the equatorial Pacific. Deep Sea Research Part II: Topical Studies in Oceanography, 49, 2601–2617.

Breitbart, M., Bonnain, C., Malki, K. & Sawaya, N. A. (2018). Phage puppet masters of the marine microbial realm. Nature Microbiology, 3, 754–766.

Brum, J. R., Morris, J. J., Décima, M. & Stukel, M. R. (2014). Mortality in the oceans: causes and consequences. Association for the Sciences of Limnology and Oceanography, 16–48.

Calbet, A. & Saiz, E. (2017). How much is enough for nutrients in microzooplankton dilution grazing experiments? Journal of Plankton Research, 40, 109–117.

Carlson, M. C. G., Ribalet, F., Maidanik, I., Durham, B. P., Hulata, Y., Ferrón, S., et al. (2022). Viruses affect picocyanobacterial abundance and biogeography in the North Pacific Ocean. Nature Microbiology, 7, 570–580.

Caron, D. A. (2017). Acknowledging and incorporating mixed nutrition into aquatic protistan ecology, finally. Environmental Microbiology Reports, 9, 41–43.

Chen, B. & Smith, S. L. (2018). CITRATE 1.0: Phytoplankton continuous trait-distribution model with one-dimensional physical transport applied to the North Pacific. Geoscientific Model Development, 11, 467–495.

Christaki, U., Giannakourou, A., Van Wambeke, F. & Grégori, G. (2001). Nanoflagellate predation on auto-and heterotrophic picoplankton in the oligotrophic Mediterranean Sea. Journal of Plankton Research, 23, 1297–1310.

Connell, P. E., Ribalet, F., Armbrust, E. V., White, A. E. & Caron, D. A. (2020). Diel oscillations in feeding strategies of heterotrophic and mixotrophic nanoplankton in the North Pacific Subtropical Gyre. Aquatic Microbial Ecology, 85, 167–181.

De Corte, D., Paredes, G., Yokokawa, T., Sintes, E. & Herndl, G. J. (2019). Differential response of *Cafeteria roenbergensis* to different Bacterial and Archaeal prey characteristics. Microbial Ecology, 78, 1–5.

Demory, D., Liu, R., Chen, Y., Zhao, F., Coenen, A. R., Zeng, Q., et al. (2020). Linking light-dependent life history traits with population dynamics for *Prochlorococcus* and cyanophage. mSystems, 5, e00586–19.

Freitas, F. H., Dugenne, M., Ribalet, F., Hynes, A., Barone, B., Karl, D. M., et al. (2020). Diel variability of bulk optical properties associated with the growth and division of small phytoplankton in the North Pacific Subtropical Gyre. Applied Optics, 59, 6702–6716.

Frias-Lopez, J., Thompson, A., Waldbauer, J. & Chisholm, S. W. (2009). Use of stable isotope-labelled cells to identify active grazers of picocyanobacteria in ocean surface waters. Environmental Microbiology, 11, 512–525.

Fuhrman, J. A. (1999). Marine viruses and their biogeochemical and ecological effects. Nature, 399, 541–548.

Fuhrman, J. A. & Noble, R. T. (1995). Viruses and protists cause similar bacterial mortality in coastal seawater. Limnology and Oceanography, 40, 1236–1242.

Ge, H., Xu, K. & Ghahramani, Z. (2018). Turing: a language for flexible probabilistic inference. In: International Conference on Artificial Intelligence and Statistics, AISTATS 2018, 9-11 April 2018, Playa Blanca, Lanzarote, Canary Islands, Spain, pp. 1682–1690.

Goldin, S., Hulata, Y., Baran, N. & Lindell, D. (2020). Quantification of T4-like and T7-like cyanophages using the polony method show they are significant members of the virioplankton in the photic zone of the North Pacific Subtropical Gyre. Frontiers in Microbiology, 11, 1210.

Grossowicz, M., Roth-Rosenberg, D., Aharonovich, D., Silverman, J., Follows, M. J. & Sher, D. (2017). *Prochlorococcus* in the lab and *in silico*: the importance of representing exudation. Limnology and Oceanography, 62, 818–835.

Groussman, R. D., Coesel, S. N., Durham, B. P. & Armbrust, E. V. (2021). Diel-Regulated Transcriptional Cascades of Microbial Eukaryotes in the North Pacific Subtropical Gyre. Frontiers in Microbiology, 12, 682651.

Gutenkunst, R. N., Waterfall, J. J., Casey, F. P., Brown, K. S., Myers, C. R. & Sethna, J. P. (2007). Universally sloppy parameter sensitivities in systems biology models. PLoS Computational Biology, 3, e189.

Hoffman, M. D., Gelman, A., et al. (2014). The No-U-Turn sampler: adaptively setting path lengths in Hamiltonian Monte Carlo. J. Mach. Learn. Res., 15, 1593–1623.

Hu, S. K., Connell, P. E., Mesrop, L. Y. & Caron, D. A. (2018). A hard day’s night: Diel shifts in microbial eukaryotic activity in the North Pacific Subtropical Gyre. Frontiers in Marine Science, 5, 351.

Hunter-Cevera, K. R., Neubert, M. G., Olson, R. J., Shalapyonok, A., Solow, A. R. & Sosik, H. M. (2020). Seasons of *Syn*. Limnology and Oceanography, 65, 1085–1102.

Hunter-Cevera, K. R., Neubert, M. G., Solow, A. R., Olson, R. J., Shalapyonok, A. & Sosik, H. M. (2014). Diel size distributions reveal seasonal growth dynamics of a coastal phytoplankter. Proceedings of the National Academy of Sciences, 111, 9852–9857.

Hynes, A. M., Rhodes, K. L. & Binder, B. J. (2015). Assessing cell cycle-based methods of measuring *Prochlorococcus* division rates using an individual-based model. Limnology and Oceanography: Methods, 13, 640–650.

Li, C., Chiang, K.-P., Laws, E. A., Liu, X., Chen, J., Huang, Y., et al. (2022). Quasi-antiphase Diel Patterns of Abundance and Cell Size/Biomass of Picophytoplankton in the Oligotrophic Ocean. Geophysical Research Letters, 49, e2022GL097753.

Li, Q., Edwards, K. F., Schvarcz, C. R., Selph, K. E. & Steward, G. F. (2021). Plasticity in the grazing ecophysiology of *Florenciella* (Dichtyochophyceae), a mixotrophic nanoflagellate that consumes *Prochlorococcus* and other bacteria. Limnology and Oceanography, 66, 47–60.

Li, Q., Edwards, K. F., Schvarcz, C. R. & Steward, G. F. (2022). Broad phylogenetic and functional diversity among mixotrophic consumers of *Prochlorococcus*. The ISME journal, 16, 1557–1569.

Liu, R., Liu, Y., Chen, Y., Zhan, Y. & Zeng, Q. (2019). Cyanobacterial viruses exhibit diurnal rhythms during infection. Proceedings of the National Academy of Sciences, 116, 14077– 14082.

Lopez, J. S., Garcia, N. S., Talmy, D. & Martiny, A. C. (2016). Diel variability in the elemental composition of the marine cyanobacterium *Synechococcus*. Journal of Plankton Research, 38, 1052–1061.

Malerba, M. E., Marshall, D. J., Palacios, M. M., Raven, J. A. & Beardall, J. (2021). Cell size influences inorganic carbon acquisition in artificially selected phytoplankton. New Phytologist, 229, 2647–2659.

Mateus, M. D. (2017). Bridging the Gap between Knowing and Modeling Viruses in Marine Systems–An Upcoming Frontier. Frontiers in Marine Science, 3, 284.

Mattern, J. P., Glauninger, K., Britten, G. L., Casey, J. R., Hyun, S., Wu, Z., et al. (2022). A Bayesian approach to modeling phytoplankton population dynamics from size distribution time series. PLoS Computational Biology, 18, e1009733.

Matteson, A. R., Rowe, J. M., Ponsero, A. J., Pimentel, T. M., Boyd, P. W. & Wilhelm, S. W. (2013). High abundances of cyanomyoviruses in marine ecosystems demonstrate ecological relevance. FEMS microbiology ecology, 84, 223–234.

Mojica, K. D., Huisman, J., Wilhelm, S. W. & Brussaard, C. P. (2016). Latitudinal variation in virus-induced mortality of phytoplankton across the North Atlantic Ocean. The ISME Journal, 10, 500–513.

Mruwat, N., Carlson, M. C. G., Goldin, S., Ribalet, F., Kirzner, S., Hulata, Y., et al. (2021). A single-cell polony method reveals low levels of infected *Prochlorococcus* in oligotrophic waters despite high cyanophage abundances. The ISME Journal, 15, 41–54.

Muratore, D., Boysen, A. K., Harke, M. J., Becker, K. W., Casey, J. R., Coesel, S. N., et al. (2022). Complex marine microbial communities partition metabolism of scarce resources over the diel cycle. Nature Ecology & Evolution, 6, 218–229.

Ng, W. H. A. & Liu, H. (2016). Diel periodicity of grazing by heterotrophic nanoflagellates influenced by prey cell properties and intrinsic grazing rhythm. Journal of Plankton Research, 38, 636–651.

Ni, T. & Zeng, Q. (2016). Diel infection of cyanobacteria by cyanophages. Frontiers in Marine Science, 2, 123.

Partensky, F., Hess, W. R. & Vaulot, D. (1999). *Prochlorococcus*, a marine photosynthetic prokaryote of global significance. Microbiology and Molecular Biology Reviews, 63, 106– 127.

Pasulka, A. L., Samo, T. J. & Landry, M. R. (2015). Grazer and viral impacts on microbial growth and mortality in the southern California Current Ecosystem. Journal of Plankton Research, 37, 320–336.

Ribalet, F., Berthiaume, C., Hynes, A., Jarred, S., Carlson, M., Clayton, S., et al. (2019a). SeaFlow data v1, high-resolution abundance, size and biomass of small phytoplankton in the North Pacific. Scientific Data, 6, 1–8.

Ribalet, F., Berthiaume, C., Hynes, A., Jarred, S., Carlson, M., Clayton, S., et al. (2019b). SeaFlow data v1: High-resolution abundance, size and biomass of small phytoplankton in the North Pacific. Version 1.2. Zenodo, doi: 10.5281/zenodo.3445407.

Ribalet, F., Swalwell, J., Clayton, S., Jiménez, V., Sudek, S., Lin, Y., et al. (2015). Light-driven synchrony of *Prochlorococcus* growth and mortality in the subtropical Pacific gyre. Proceedings of the National Academy of Sciences, 112, 8008–8012.

Sanders, R. W. & Porter, K. G. (1988). Phagotrophic phytoflagellates. In: Advances in microbial ecology (ed.). Springer, pp. 167–192.

Sherr, E. B., Caron, D. A. & Sherr, B. F. (1993). Staining of heterotrophic protists for visualization via epifluorescence microscopy. Lewis Publishers, Boca Raton, pp. 213–227.

Sosik, H. M., Olson, R. J., Neubert, M. G., Shalapyonok, A. & Solow, A. R. (2003). Growth rates of coastal phytoplankton from time-series measurements with a submersible flow cytometer. Limnology and Oceanography, 48, 1756–1765.

Stoecker, D. K., Hansen, P. J., Caron, D. A. & Mitra, A. (2017). Mixotrophy in the marine plankton. Annu. Rev. Mar. Sci, 9, 311–335.

Sullivan, M. B., Weitz, J. S. & Wilhelm, S. (2017). Viral ecology comes of age. Environmental Microbiology Reports, 9, 33–35.

Talmy, D., Beckett, S. J., Taniguchi, D. A. A., Brussaard, C. P., Weitz, J. S. & Follows, M. J. (2019a). An empirical model of carbon transfer to marine viruses and zooplankton grazers. Environmental Microbiology, 21, 2171–2181.

Talmy, D., Beckett, S. J., Zhang, A. B., Taniguchi, D. A. A., Weitz, J. S. & Follows, M. J. (2019b). Contrasting controls on microzooplankton grazing and viral infection of microbial prey. Frontiers in Marine Science, 6, 182.

Taniguchi, D. A. A., Franks, P. J. & Poulin, F. J. (2014). Planktonic biomass size spectra: an emergent property of size-dependent physiological rates, food web dynamics, and nutrient regimes. Marine Ecology Progress Series, 514, 13–33.

Thamatrakoln, K., Talmy, D., Haramaty, L., Maniscalco, C., Latham, J. R., Knowles, B., et al. (2018). Light regulation of coccolithophore host–virus interactions. New Phytologist, 221, 1289–1302.

Tsakalakis, I., Follows, M. J., Dutkiewicz, S., Follett, C. L. & Vallino, J. J. (2022). Diel light cycles affect phytoplankton competition in the global ocean. Global Ecology and Biogeography, 31, 1838–1849.

Vaulot, D. (1995). The cell cycle of phytoplankton: coupling cell growth to population growth. In: Molecular ecology of aquatic microbes. Springer, pp. 303–322.

Vaulot, D. & Marie, D. (1999). Diel variability of photosynthetic picoplankton in the equatorial Pacific. Journal of Geophysical Research: Oceans, 104, 3297–3310.

Vislova, A., Sosa, O. A., Eppley, J. M., Romano, A. E. & DeLong, E. F. (2019). Diel oscillation of microbial gene transcripts declines with depth in oligotrophic ocean waters. Frontiers in Microbiology, 10, 2191.

Weinbauer, M. G. (2004). Ecology of prokaryotic viruses. FEMS Microbiology Reviews, 28, 127–181.

Weitz, J. S., Stock, C. A., Wilhelm, S. W., Bourouiba, L., Coleman, M. L., Buchan, A., et al. (2015). A multitrophic model to quantify the effects of marine viruses on microbial food webs and ecosystem processes. The ISME Journal, 9, 1352.

Weitz, J. S. & Wilhelm, S. W. (2012). Ocean viruses and their effects on microbial communities and biogeochemical cycles. F1000 biology reports, 4.

Welkie, D. G., Rubin, B. E., Diamond, S., Hood, R. D., Savage, D. F. & Golden, S. S. (2018). A Hard Day’s Night: Cyanobacteria in Diel Cycles. Trends in Microbiology, 27, 231–242.

Wilhelm, S. W. & Suttle, C. A. (1999). Viruses and nutrient cycles in the sea: viruses play critical roles in the structure and function of aquatic food webs. Bioscience, 49, 781–788.

Wirtz, K. W. (2019). Physics or biology? Persistent chlorophyll accumulation in a shallow coastal sea explained by pathogens and carnivorous grazing. PLOS ONE, 14, e0212143.

Wu, Z., Aharonovich, D., Roth-Rosenberg, D., Weissberg, O., Luzzatto-Knaan, T., Vogts, A., et al. (2022). Single-cell measurements and modelling reveal substantial organic carbon acquisition by *Prochlorococcus*. Nature Microbiology, 7, 2068–2077.

## References

Baines, S. B., Twining, B. S., Brzezinski, M. A., Krause, J. W., Vogt, S., Assael, D., et al. (2012). Significant silicon accumulation by marine picocyanobacteria. Nature Geoscience, 5, 886–891.

Baker, K. G. & Geider, R. J. (2021). Phytoplankton mortality in a changing thermal seascape. Global Change Biology, 27, 5253–5261.

Berg, H. C. & Purcell, E. M. (1977). Physics of chemoreception. Biophysical Journal, 20, 193– 219.

Bidle, K. D. (2015). The molecular ecophysiology of programmed cell death in marine phytoplankton. Annual review of marine science, 7, 341–375.

Bidle, K. D. (2016). Programmed cell death in unicellular phytoplankton. Current Biology, 26, R594–R607.

Biller, S. J., Berube, P. M., Lindell, D. & Chisholm, S. W. (2015). *Prochlorococcus*: the structure and function of collective diversity. Nature Reviews Microbiology, 13, 13.

Cai, L., Chen, Y., Xiao, S., Liu, R., He, M., Zhang, R., et al. (2022). Abundant and cosmopolitan lineage of cyanopodoviruses lacking a DNA polymerase gene. The ISME Journal, 1–11.

Casey, J. R., Björkman, K. M., Ferrón, S. & Karl, D. M. (2019). Size dependence of metabolism within marine picoplankton populations. Limnology and Oceanography, 64, 1819–1827.

Cruz, B. N. & Neuer, S. (2019). Heterotrophic bacteria enhance the aggregation of the marine picocyanobacteria *Prochlorococcus* and *Synechococcus*. Frontiers in Microbiology, 10, 1864.

Demory, D., Combe, C., Hartmann, P., Talec, A., Pruvost, E., Hamouda, R., et al. (2018). How do microalgae perceive light in a high-rate pond? Towards more realistic Lagrangian experiments. Royal Society open science, 5, 180523.

Einstein, A. (1905). Über die von der molekularkinetischen Theorie der Wärme geforderte Bewegung von in ruhenden Flüssigkeiten suspendierten Teilchen. Annalen der physik, 322, 549–560.

Franklin, D. J. (2013). Explaining the causes of cell death in cyanobacteria: what role for asymmetric division? Journal of Plankton Research, 36, 11–17.

Fucich, D. & Chen, F. (2020). Presence of toxin-antitoxin systems in picocyanobacteria and their ecological implications. The ISME Journal, 14, 2843–2850.

Gelman, A., Carlin, J. B., Stern, H. S. & Rubin, D. B. (1995). Bayesian data analysis. Chapman and Hall/CRC.

Guidi, L., Chaffron, S., Bittner, L., Eveillard, D., Larhlimi, A., Roux, S., et al. (2016). Plankton networks driving carbon export in the oligotrophic ocean. Nature, 532, 465.

Iuculano, F., Mazuecos, I. P., Reche, I. & Agustí, S. (2017). *Prochlorococcus* as a possible source for transparent exopolymer particles (TEP). Frontiers in Microbiology, 8, 709.

Johnson, Z. I., Zinser, E. R., Coe, A., McNulty, N. P., Woodward, E. M. S. & Chisholm, S. W. (2006). Niche partitioning among *Prochlorococcus* ecotypes along ocean-scale environmental gradients. Science, 311, 1737–1740.

Karayanni, H., Christaki, U., Van Wambeke, F., Thyssen, M. & Denis, M. (2008). Heterotrophic nanoflagellate and ciliate bacterivorous activity and growth in the northeast Atlantic Ocean: a seasonal mesoscale study. Aquatic Microbial Ecology, 51, 169–181.

Karl, D. M. & Church, M. J. (2014). Microbial oceanography and the Hawaii Ocean Time-series programme. Nature Reviews Microbiology, 12, 699–713.

Kearney, S. M., Thomas, E., Coe, A. & Chisholm, S. W. (2021). Microbial Diversity of Cooccurring Heterotrophs in Cultures of Marine Picocyanobacteria. Environmental Microbiome, 16, 1–15.

Kiørboe, T. (2008). A mechanistic approach to plankton ecology. Princeton University Press.

Kiørboe, T. (2011). How zooplankton feed: mechanisms, traits and trade-offs. Biological Reviews, 86, 311–339.

Krause, J. W., Brzezinski, M. A., Baines, S. B., Collier, J. L., Twining, B. S. & Ohnemus, D. C. (2017). Picoplankton contribution to biogenic silica stocks and production rates in the Sargasso Sea. Global Biogeochemical Cycles, 31, 762–774.

Kulk, G., Poll, W. H. van de, Visser, R. J. & Buma, A. G. (2013). Low nutrient availability reduces high-irradiance-induced viability loss in oceanic phytoplankton. Limnology and Oceanography, 58, 1747–1760.

Lambert, B. (2018). A student’s guide to Bayesian statistics. Sage.

Li, Q., Edwards, K. F., Schvarcz, C. R., Selph, K. E. & Steward, G. F. (2021). Plasticity in the grazing ecophysiology of *Florenciella* (Dichtyochophyceae), a mixotrophic nanoflagellate that consumes *Prochlorococcus* and other bacteria. Limnology and Oceanography, 66, 47– 60.

Link, W. A. & Eaton, M. J. (2012). On thinning of chains in MCMC. Methods in ecology and evolution, 3, 112–115.

Llabrés, M., Agustí, S. & Herndl, G. J. (2011). Diel in situ picophytoplankton cell death cycles coupled with cell division. Journal of phycology, 47, 1247–1257.

Mann, E. L., Ahlgren, N., Moffett, J. W. & Chisholm, S. W. (2002). Copper toxicity and cyanobacteria ecology in the Sargasso Sea. Limnology and Oceanography, 47, 976–988.

Mann, E. L. & Chisholm, S. W. (2000). Iron limits the cell division rate of *Prochlorococcus* in the eastern equatorial Pacific. Limnology and Oceanography, 45, 1067–1076.

Martiny, A. C., Coleman, M. L. & Chisholm, S. W. (2006). Phosphate acquisition genes in *Prochlorococcus* ecotypes: evidence for genome-wide adaptation. Proceedings of the National Academy of Sciences, 103, 12552–12557.

Mella-Flores, D., Six, C., Ratin, M., Partensky, F., Boutte, C., Le Corguillé, G., et al. (2012). *Prochlorococcus* and *Synechococcus* have evolved different adaptive mechanisms to cope with light and UV stress. Frontiers in Microbiology, 3, 285.

Menden-Deuer, S. & Lessard, E. J. (2000). Carbon to volume relationships for dinoflagellates, diatoms, and other protist plankton. Limnology and Oceanography, 45, 569–579.

Moger-Reischer, R. Z. & Lennon, J. T. (2019). Microbial ageing and longevity. Nature Reviews Microbiology, 17, 679–690.

Moore, L. R. & Chisholm, S. W. (1999). Photophysiology of the marine cyanobacterium *Prochlorococcus*: ecotypic differences among cultured isolates. Limnology and Oceanography, 44, 628–638.

Morris, J. J., Johnson, Z. I., Szul, M. J., Keller, M. & Zinser, E. R. (2011). Dependence of the cyanobacterium *Prochlorococcus* on hydrogen peroxide scavenging microbes for growth at the ocean’s surface. PLOS ONE, 6, e16805.

Morris, J. J., Kirkegaard, R., Szul, M. J., Johnson, Z. I. & Zinser, E. R. (2008). Facilitation of robust growth of *Prochlorococcus* colonies and dilute liquid cultures by “helper” heterotrophic bacteria. Applied and Environmental Microbiology, 74, 4530–4534.

Murata, K., Zhang, Q., Galaz-Montoya, J. G., Fu, C., Coleman, M. L., Osburne, M. S., et al. (2017). Visualizing adsorption of cyanophage P-SSP7 onto marine *Prochlorococcus*. Scientific Reports, 7, 44176.

Murray, A. G. & Jackson, G. A. (1992). Viral dynamics: a model of the effects of size, shape, motion and abundance of single-celled planktonic organisms and other particles. Marine Ecology Progress Series, 103–116.

Neuer, S. & Cowles, T. J. (1994). Protist herbivory in the Oregon upwelling system. Marine ecology progress series., 113, 147–162.

Pittera, J., Humily, F., Thorel, M., Grulois, D., Garczarek, L. & Six, C. (2014). Connecting thermal physiology and latitudinal niche partitioning in marine *Synechococcus*. The ISME journal, 8, 1221–1236.

Ribalet, F., Berthiaume, C., Hynes, A., Jarred, S., Carlson, M., Clayton, S., et al. (2019). SeaFlow data v1, high-resolution abundance, size and biomass of small phytoplankton in the North Pacific. Scientific Data, 6, 1–8.

Richardson, T. L. (2019). Mechanisms and pathways of small-phytoplankton export from the surface ocean. Ann. Rev. Mar. Sci, 11, 57–74.

Richardson, T. L. & Jackson, G. A. (2007). Small phytoplankton and carbon export from the surface ocean. Science, 315, 838–840.

Roth-Rosenberg, D., Aharonovich, D., Luzzatto-Knaan, T., Vogts, A., Zoccarato, L., Eigemann, F., et al. (2020). *Prochlorococcus* Cells Rely on Microbial Interactions Rather than on Chlorotic Resting Stages To Survive Long-Term Nutrient Starvation. mBio, 11, e01846– 20.

Ruf, T. (1999). The Lomb-Scargle Periodogram in Biological Rhythm Research: Analysis of Incomplete and Unequally Spaced Time-Series. Biological Rhythm Research, 30, 178–201.

Sarker, I., Moore, L. R. & Tetu, S. G. (2021). Investigating zinc toxicity responses in marine *Prochlorococcus* and *Synechococcus*. Microbiology, 167, 001064.

Straile, D. (1997). Gross growth efficiencies of protozoan and metazoan zooplankton and their dependence on food concentration, predator-prey weight ratio, and taxonomic group. Limnology and Oceanography, 42, 1375–1385.

Stramska, M. & Dickey, T. D. (1998). Short-term variability of the underwater light field in the oligotrophic ocean in response to surface waves and clouds. Deep Sea Research Part I: Oceanographic Research Papers, 45, 1393–1410.

Thaben, P. F. & Westermark, P. O. (2014). Detecting rhythms in time series with RAIN. Journal of biological rhythms, 29, 391–400.

Thompson, A. W., Huang, K., Saito, M. A. & Chisholm, S. W. (2011). Transcriptome response of high-and low-light-adapted *Prochlorococcus* strains to changing iron availability. The ISME Journal, 5, 1580–1594.

Tolonen, A. C., Aach, J., Lindell, D., Johnson, Z. I., Rector, T., Steen, R., et al. (2006). Global gene expression of *Prochlorococcus* ecotypes in response to changes in nitrogen availability. Molecular systems biology, 2, 53.

Vardi, A., Formiggini, F., Casotti, R., De Martino, A., Ribalet, F., Miralto, A., et al. (2006). A stress surveillance system based on calcium and nitric oxide in marine diatoms. PLOS Biology, 4, e60.

Verity, P. G., Stoecker, D. K., Sieracki, M. E. & Nelson, J. R. (1993). Grazing, growth and mortality of microzooplankton during the 1989 North Atlantic spring bloom at 47 N, 18 W. Deep Sea Research Part I: Oceanographic Research Papers, 40, 1793–1814.

Von Smoluchowski, M. (1917). Investigation of a mathematical theory on the coagulation of colloidal suspensions. Z. Physik. Chem.(Ger.), 92, 155.

Wei, Y., Sun, J., Chen, Z., Zhang, Z., Zhang, G. & Liu, X. (2021). Significant contribution of picoplankton size fraction to biogenic silica standing stocks in the Western Pacific Ocean. Progress in Oceanography, 192, 102516.

Xiao, X., Guo, W., Li, X., Wang, C., Chen, X., Lin, X., et al. (2021). Viral lysis alters the optical properties and biological availability of dissolved organic matter derived from picocyanobacteria *Prochlorococcus*. Applied and Environmental Microbiology, 87, e02271– 20.

Zehr, J. P., Weitz, J. S. & Joint, I. (2017). How microbes survive in the open ocean. Science, 357, 646–647.

Zhao, Z., Gonsior, M., Schmitt-Kopplin, P., Zhan, Y., Zhang, R., Jiao, N., et al. (2019). Microbial transformation of virus-induced dissolved organic matter from picocyanobacteria: coupling of bacterial diversity and DOM chemodiversity. The ISME Journal, 13, 2551– 2565.

Zheng, Q., Lin, W., Wang, Y., Li, Y., He, C., Shen, Y., et al. (2021). Highly enriched N-containing organic molecules of *Synechococcus* lysates and their rapid transformation by heterotrophic bacteria. Limnology and Oceanography, 66, 335–348.

Zinser, E. R., Johnson, Z. I., Coe, A., Karaca, E., Veneziano, D. & Chisholm, S. W. (2007). Influence of light and temperature on *Prochlorococcus* ecotype distributions in the Atlantic Ocean. Limnology and Oceanography, 52, 2205–2220.

Zubkov, M. V. & Tarran, G. A. (2008). High bacterivory by the smallest phytoplankton in the North Atlantic Ocean. Nature, 455, 224–226.

